# Combined online Bayesian and windowed estimation of background and signal localization facilitates active-feedback particle tracking in complex environments

**DOI:** 10.1101/2022.08.05.502973

**Authors:** Anastasia Niver, Kevin D. Welsher

## Abstract

Despite successes in tracking single molecules *in vitro*, the extension of active-feedback single-particle methods to tracking rapidly diffusing and unconfined proteins in live cells has not been realized. Since existing active-feedback localization methods localize particles in real time assuming zero background, they are ill-suited to track in the inhomogeneous background environment of a live cell. Here, we develop a windowed estimation of signal and background levels which uses recent data to estimate the current particle brightness and background intensity. These estimates facilitate recursive Bayesian position estimation, improving upon current Kalman-based localization methods. Combined, online Bayesian and windowed estimation of background and signal (COBWEBS) surpasses existing localization methods. Simulations demonstrate improved localization accuracy and responsivity in a homogenous background for selected particle and background intensity combinations. Improved or similar performance of COBWEBS tracking extends to the majority of signal and background combinations explored. Furthermore, improved tracking durations are demonstrated in the presence of heterogeneous backgrounds for multiple particle intensities, diffusive speeds, and background patterns. COBWEBS can accurately track particles in the presence of high and non-uniform backgrounds including intensity changes of up to three-fold that of the particle’s intensity, making it a prime candidate for advancing active-feedback single-fluorophore tracking to the cellular interior.

## I. Introduction

Active-feedback particle tracking microscopy applies to an increasing range of critical biological and chemical research questions.^1–9^ Despite successes in tracking single dye labeled molecules *in vitro* and brighter particles in live cells, the extension of these powerful methods to tracking rapidly diffusing and unconfined proteins in live cells has not been realized. Tetrahedral approaches have been applied to single fluorescent dyes and fluorescent proteins in glycerol/water solutions, as well as to membrane bound or vesicle confined quantum dot (bright) labeled proteins in live cells.^10, 11^ Orbital tracking has been applied to study mitochondrial trafficking along vertebrate axons and artificial virus trafficking along microtubules in vivo, both of which are bright particles with many fluorescent labels.^12, 13^ Recently, 3D single-molecule active real-time tracking (3D-SMART) has been applied to track single fluorophore-labelled biomolecules (including DNA, proteins, and RNA) in glycerol/water solutions, the binding of fluorescent protein labelled lentiviral particles to live cells from the extracellular space, protein corona formation on nanoparticles, and the chemical kinetics of single polymer particles.^14–18^ Collectively, single-particle active-feedback tracking techniques successfully follow single fluorophores in vitro but require multiply labeled bright slow-moving particles for tracking in live cells.

Live cells present a challenging tracking environment. Even in the absence of non-specific labeling or spectral bleed-through, the cell itself generates unavoidable background fluorescence which originates from biomolecules including FAD and NADH.^19^ This autofluorescence shows clustering distributed throughout the cytoplasm and cannot always be filtered out.^20^ Lifetime gating has shown some promise in this area, but relies on emitters with longer lifetimes than the background such as quantum dots.^21^ To translate real-time single-fluorophore tracking to the cellular interior, improving upon how each of these active-feedback methods deal with background photons is critical.

To date, active-feedback tracking experiments have relied on relatively simple algorithms to estimate particle positions. The simplicity is necessary as the molecule must be localized and feedback applied in real time before the molecule has a chance to diffuse out of the detection volume, roughly on the millisecond timescale. Increased algorithm complexity comes at the cost of increased position estimate computation time, limiting the responsivity. To maintain estimate computation times below the physical response speed, most tracking implementations use intensity changes to detect shifts in particle positions.^9, 11–13^ The 3D-SMART approach localizes particles using an assumed Gaussian recursive estimation which weighs more recent photon arrivals more heavily but still localizes the particle to the area of highest intensity.^14^ The zero-background assumption of existing methods interferes with particle position estimations when intense and non-homogeneous backgrounds are present. In this work, we demonstrate a novel windowed signal and background estimation algorithm. Utilizing a subset of the most recent photons, we estimate intensity of both the tracked particle and its environment. These signal and background estimates facilitate position estimation using a recursive Bayesian algorithm. The combined online Bayesian and windowed estimation of background and signal (COBWEBS) method can accurately track particles in the presence of high and non-uniform background. The COBWEBS algorithm presented herein is robust to large and abruptly changing signal-to-background ratios, making it a prime candidate for advancing active feedback single fluorophore tracking to the cellular interior.

## II. Methods

### A. Kalman

An assumed-Gaussian density filter mapped onto a Kalman filter suitable for position estimation is fully derived in Fields and Cohen.^22^ The key results are summarized below. Several assumptions go into this derivation. The point spread function (PSF) of the system is assumed to be a symmetric Gaussian function. Additionally, the laser spot covariance, *W*, is assumed to be diagonal which is the case as long as beam asymmetry is along the same axes as feedback is applied.^22^ There are two major assumptions that allow all likelihoods to be modelled as Gaussians. First, it is assumed that the background level is zero. In practice, this means that each arriving photon is assumed to originate from the tracked molecule, which in very low background environments is generally true. Second, the laser scan rate is assumed to be high relative to the rate of photon detection. In other words, each bin within the scan will contain one photon at most. After these assumptions the update equations from an assumed Gaussian density filter simplify to Eq. 1 and Eq. 2. The current particle position is a function of laser covariance *W*, number of photons detected *n*, and uncertainty associated with the previous position estimate *σ*. The subscripts *k* and *k* – 1 assign a value to either the current bin or the previous bin. The position estimate given by the calculated mean of the assumed gaussian position estimate is *x*. The laser spot center position is *c*.

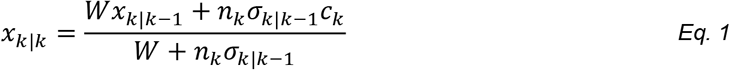

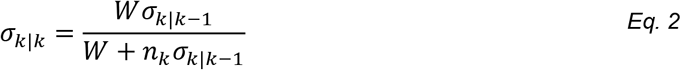

Both the particle and stage can be treated as stationary during position estimation since their step sizes are on the order of 10^-8^ m (for D ~ 1 to 10 μm^2^/s particles) during a 20 μs bin, orders of magnitude smaller than the laser scan area of 10^-6^ m. The stage motions applied during each bin to correct for particle diffusion are of a similarly negligible magnitude. This simplifies the prediction step of the Kalman filter to Eq. 3 and Eq. 4. where *x* and *σ* remain as defined above, *D* is the diffusion coefficient, and tau is the bin time of a given estimate. This model assumes isotropic random diffusion.

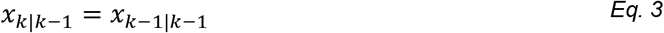

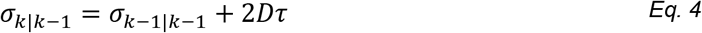

The estimated particle position, *x_k_*, is fed into a proportional-integral-derivative (PID) feedback controller (Eq. S2) which drives the response mechanism of the feedback tracking method and recenters the particle. The simplicity of this Kalman filter makes it ideal for numerical implementation on a field-programmable gate array (FPGA). The filter works extremely well for 3D-SMART in low background environments, but the baked-in assumption of zero background has dramatic consequences for tracking in realistic environments.

### B. Combined online Bayesian and windowed estimation of background and signal (COBWEBS)

#### 1. Online Bayesian

To develop a position estimation algorithm that can account for non-zero and non-uniform background, we apply recursive Bayesian inference and remove the assumption that all likelihoods can be approximated as Gaussian, as used in Kalman estimation. Assuming a laser excitation power below saturation, the rate of expected photon detection *γ_k_* for a given bin can be modeled as a Gaussian function of the distance between the particle and the laser spot, plus a background emission rate.^22–24^ Here, *c_k_* is the laser spot center position for a given bin, *s* is the maximum emitter (signal) count rate in counts per second (cps) for a given bin, and *b*(*x, y*) is the non-emitter (background) count rate in cps described as a function of position since the background is spatially dependent. Assuming that the covariance matrix is diagonal, similar to the Kalman case above, this calculation becomes Eq. 5 where *σ* is the width of the point-spread function (PSF) and *x* and *y* are the particle’s position along a given axis.

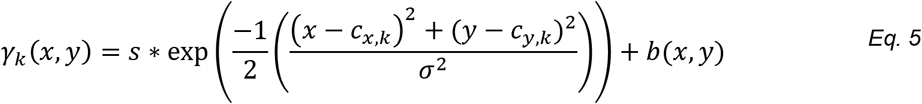

The probability of detecting *n* photons in a bin of time *τ* is given by a Poisson distribution (Eq. 6).

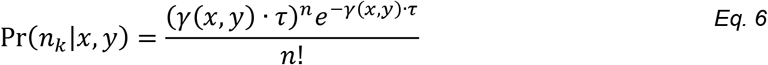

The update step combines the probability of detecting *n* photons given a particle at position (*x_k_, y_k_*) with the prior probability of finding a particle at that position.

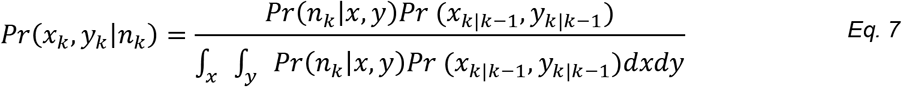

Here, Pr(*x_k_, y_k_|n_k_*) is the normalized posterior likelihood distribution and *Pr* (*x*_*k|k*-1_, *y*_*k|k*-1_) is the prior likelihood distribution given all previous observations. The denominator on the RHS of Eq. 7 is simply a normalization factor. The maximum of the posterior distribution yields a maximum-likelihood estimate (MLE) of the current particle position. For tracking, this MLE is fed into a proportional-integral-derivative (PID) feedback controller (Eq. S2).

For a stationary particle, this posterior distribution would become the prior distribution for the next observation bin and the process would continue. However, for a moving particle, uncertainty introduced by particle motion must be accounted for. Here, we use numerical convolution of the expected diffusion kernel (Eq. S1) with the posterior distribution from Eq. 7 to incorporate uncertainty in particle location due to diffusion. The output of this convolution then becomes the prior distribution for the next observation bin and the process continues.

For simulations below, this analytical form of recursive Bayesian localization was converted to a numerical implementation (SI Note 1.4). Additionally, computational simplifications were developed to both improve computation speeds and ensure plausibility of the eventual experimental implementation of COBWEBS localization using a field-programmable gate array (FPGA) (SI Note 1.5). An alternative computationally simplified Bayesian implementation of COBWEBS, COBWEBS-1D, was developed by assuming that the probability distribution of particle positions along the X and Y axes was entirely independent (SI Note 2). Assumed independence reduces the number of elements in the prior and posterior distributions from *V*^2^ to 2*V* where *V* is the number of elements along a given axis (assuming equivalent grid spacing). The windowed estimation of signal and background discussed below was identical between both approaches.

#### 2. Windowed Estimation of Background and Signal

The Bayesian position estimation described above requires knowledge of a particle’s signal (*s*) and background (*b*) (Eq. 5). A priori selection of *s* and *b* values would introduce potential bias in particle selection and preclude responsivity to changing intensities, such as photobleaching or diffusing near a brighter background object. Therefore, online estimation of *s* and *b* is critical. These estimates must replace the true values of *s* and *b*, which cannot be experimentally identified in real-time, to enable recursive Bayesian estimation of particle positions. The laser scan in 3D-SMART uses a 5×5 knights tour pattern (Figure S3 Inset) which covers a 1 μm square.^25, 26^ Since the scan area is larger than the particle’s PSF (σ = 0.15 μm), and a well tracked particle remains near the center of the scan, the proportion of the observed signal originating from the actual particle varies across the scan area. The variable contribution of *s* and *b* enables estimation of *s* and *b* values from the observed count rate (OCR) at a given scan position. Since photon arrivals are noisy, these OCRs are calculated over a sliding window using the most recent photons to generate *s* and *b* estimates which adapt to changing conditions over time. An OCR can also be calculated over the entire scan area and used to approximate experimentally observed particle intensities for a range of *s* values (Figure S3).

### C. Simulation Implementation

Particle diffusion, photon generation, and stage response were modeled using pseudo random number generation and previously established numerical methods (SI Note 1).

## III. Results

### A. Signal and background estimation

Prior to implementation of the full algorithm, the dependence of the OCR at each pixel on the true value of *s* and *b* need to be established. Deviations between the center of the stage/laser scan and the actual particle position during a typical tracking experiment were approximated as a normal distribution with a standard deviation of 0.1 μm. Pseudo random particle positions were generated from this normal approximation, then photons were generated for a range of signal and background values. Trials over a range of signal values at zero background (Fig. 2A) and vice versa (Fig. 2b) were used to establish the relationship between true *s* and *b* values and the OCR for a given pixel type. For combinations of *s* and *b* values where both values are non-zero, the OCR at the corner pixels (F) was found to be 0.003*s* + *b*. The OCR at the central pixel (A) was 0.69*s* + *b*. These expressions form a solvable system of linear equations used to extract estimates for *s* and *b* from the OCR (Figure *2*). This pair of A and B pixels was found to be the most accurate combination for signal and background estimation from the six available pixel types (Figure S5).

**Figure 1.**
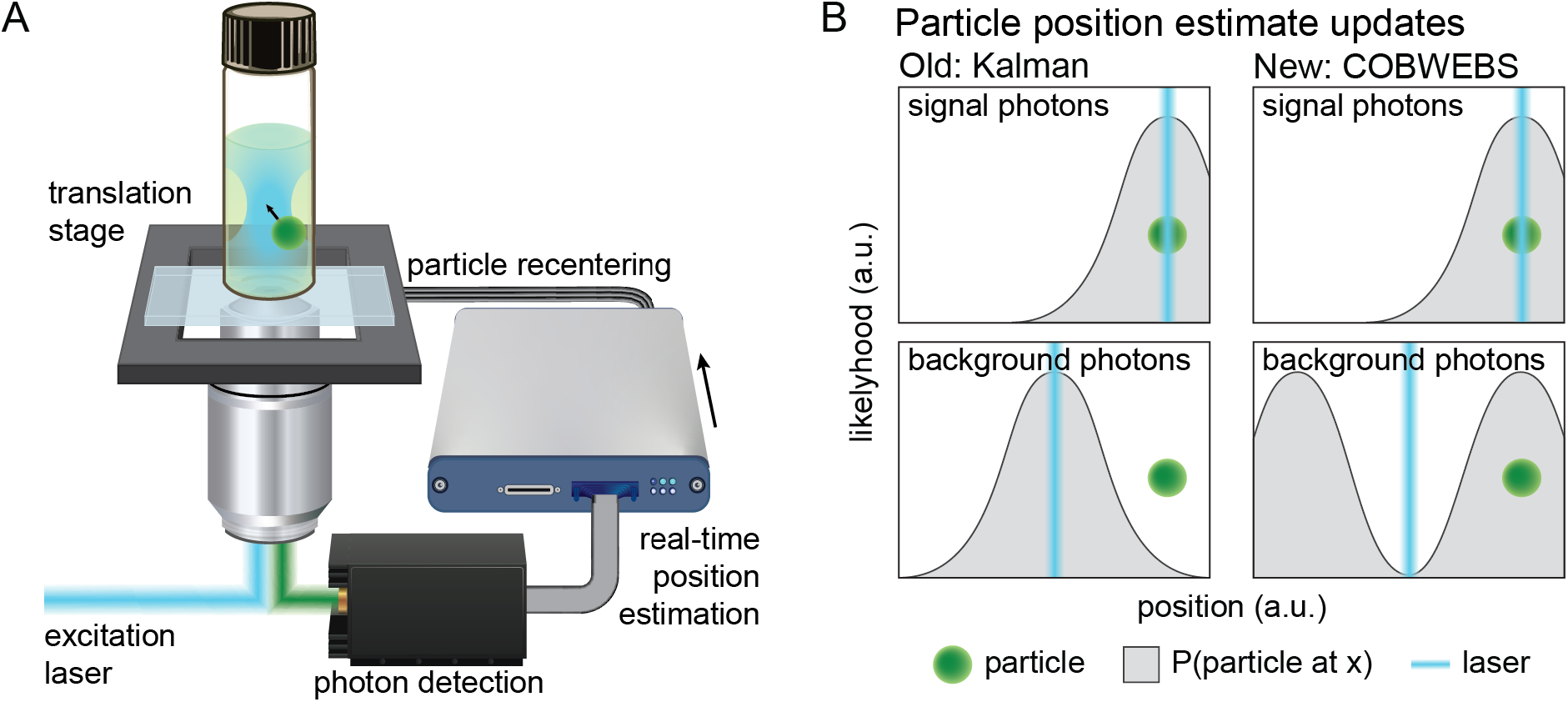
Principles of active-feedback particle tracking and position estimation algorithms. **A.** General schematic of active-feedback single-particle tracking. The excitation laser is scanned through a search pattern. Fluorescent photons are filtered and detected. Counted photons and their arrival positions are used to calculate an estimate of the particle’s position. Then, the position estimate is fed into the translation stage which is moved to recenter the tracked particle in the search area. **B.** Probability distributions of particle positions used in estimate update steps after photon detection for both the prior Kalman and new COBWEBS methods.

**Figure 2.**
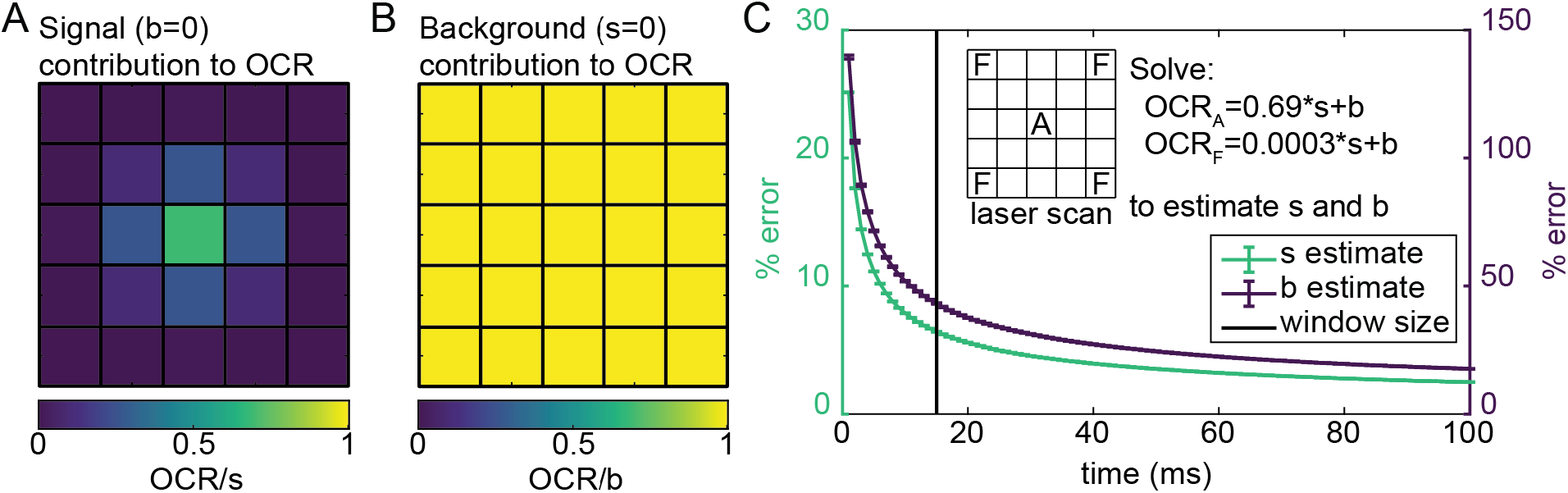
Development of real time signal and background estimation for recursive Bayesian localization. **A.** Ratio of OCR to input *s* value ratios over a range of *s* values with *b* = 0 (500 trials of 100 ms each). **B.** Ratio of OCR to input *b* values with *s* = 0. **C**. Percent error in *s* and *b* estimates vs window size used to calculate OCRs. Error bars are the standard error of the mean. The vertical line at 15 ms indicates the window selected for use in the simulations explored below. Inset: equations and pixels used to calculate estimates.

Next, the time over which the values of *s* and *b* are extracted from the OCR was determined. The time window used to determine the OCR for a given pixel type was varied from 1 to 100 ms. Larger time windows for calculation of OCRs have more accurate *s* and *b* estimation (Figure *2*C). A window width of the most recent 15 ms worth of data to calculate OCRs was selected for subsequent simulations. Although using windows larger than 15 ms can further reduce the percent error in *s* and *b* estimates by approximately a factor of two (Figure *2*C), this reduces the potential responsivity of the estimates to environmental changes.

### B. Particle tracking in uniform background environments

Performance of the Kalman and COBWEBS methods were tested first in uniform background environments. Three different classes of particles were tested, each with a different signal level. These were dim particles (overall OCR of 19 kcps), particles with emission similar to virus-like particles (VLPs, 47 kcps) and bright particles (93 kcps). We note here that the overall OCR for each class of particle is not the value s, but is reduced compared to this theoretical maximum. In practice the actual OCR will vary with tracking performance due to changes in the offset between particle position and scan center. An approximation of this conversion using the signal and background estimation data with OCR equal to 0.09043*s* was used here (Figure S3).

The first set of simulations compares simulated tracking of a VLP in an even background environment. A simulated particle with an OCR of 47 kcps was allowed to diffuse through a high background of 50 kcps (Figure *3*A). Each simulated trajectory outputs particle positions, particle position estimates, and the resulting simulated stage positions (example segments in Figure *3*B). Qualitatively, there is more scatter in the Kalman estimation than is present in the example COBWEBS position estimates for the same sequence of particle positions (Figure *3*B).

**Figure 3.**
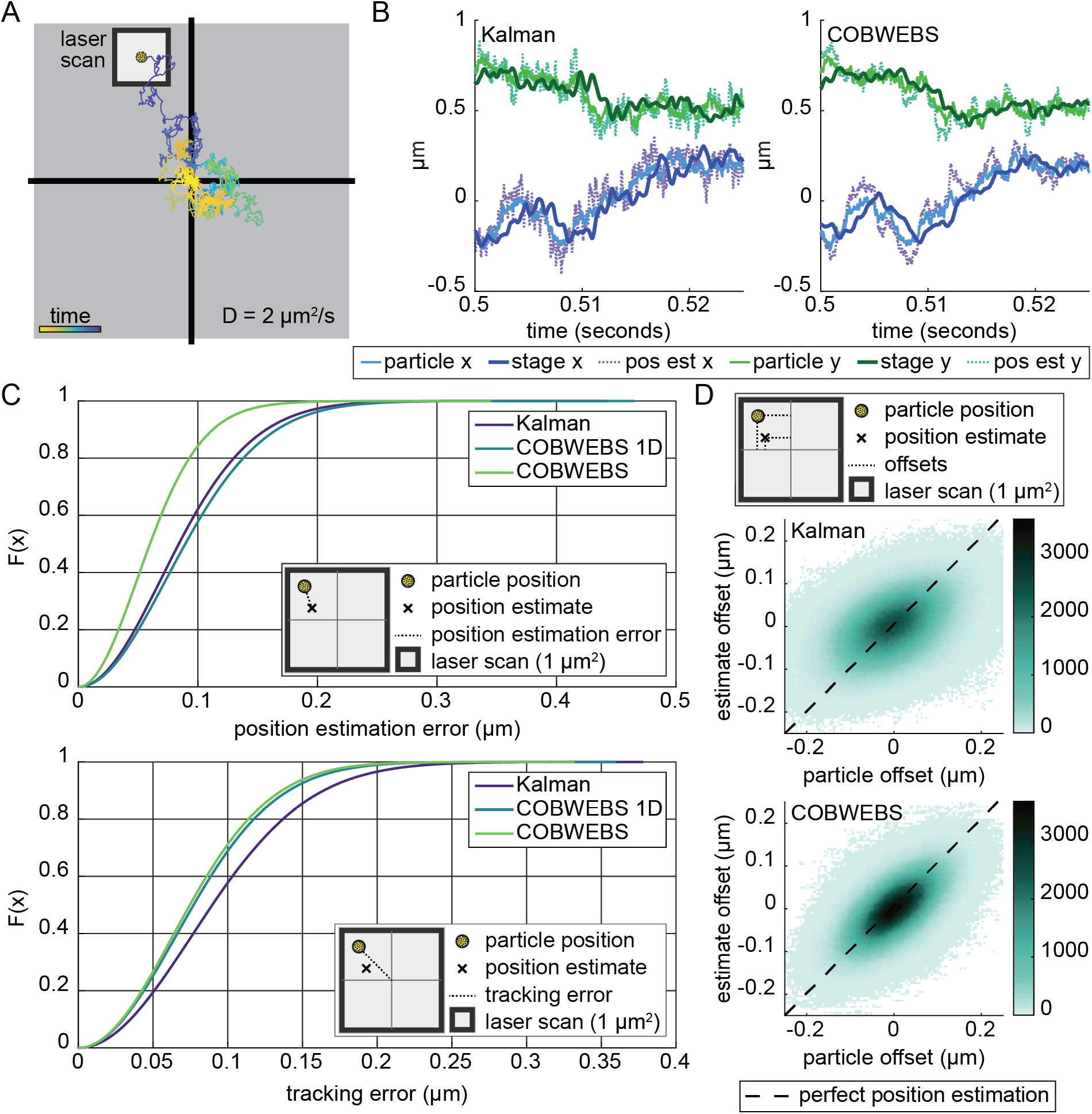
Direct comparison of error between position estimation algorithms for a single particle in an even background. **A.** Cartoon of tracking in an even background. **B.** Segments of example trajectories from Kalman and COBWEBS tracking showing the actual particle positions, the estimated particle positions, and the resulting simulated stage response. **C.** Empirical cumulative density function plots of both the position estimation error and the tracking error of each observed error value in 25 one second trajectories for all three tracking algorithms (s = 516.64 kcps, *b* = 50 kcps, D = 2 μm^2^/s). **D.** 2D histogram of particle position estimate offsets vs the physical particle offsets. The correlation coefficients for Kalman, and 2D Bayesian estimation are 0.4501 and 0.6325 respectively.

Algorithm tracking performance is compared using two types of error. Tracking error is the offset between the particle and the stage positions. Position estimation error is the difference between the estimated particle position and the particle’s true position. These errors are calculated using Eq. 8 and Eq. 9 respectively. *Stg* is the stage position along a given axis, *P* is the particle position along a given axis, and *P_Est_*. represents the estimated position along a given axis.

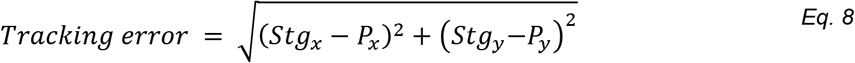

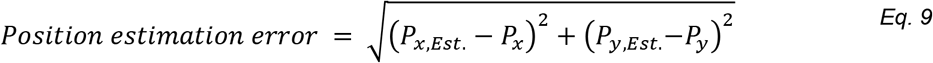

In a high background, COBWEBS tracking minimizes both tracking and position estimation error (Figure *3*C). The difference between the mean error of each trial for all three algorithms (Kalman, 1D-COBWEBS, and 2D COBWEBS) for both types of error was statistically significant (one-way ANOVA p≤2e-52 followed by post hoc Tukey-Kramer test, p≤1E-08, population of mean error value for 25 trajectories, details in Table S1 and S2).

In addition to reducing errors, COBWEBS tracking has higher responsivity to particle motions than Kalman estimation. A 2D histogram of particle displacement from the center of the laser scan vs the displacement in the corresponding position estimate illustrates this responsivity (Figure *3*D). A perfect estimator where estimated position and actual position are equal would have a correlation coefficient of one indicating a perfect linear relationship between the estimated and actual position. Moving from Kalman estimation to COBWEBS increases the correlation coefficient from 0.45 to 0.63. The high correlation for COBWEBS is consistent with its status as the best performing algorithm among the three.

Running uniform background tracking simulations for a range of signal and background levels shows COBWEBS tracking stability over a similar or improved range compared to Kalman tracking (Figure S7). Additionally, the minimum position estimation error is lower for COBWEBS tracking than the corresponding Kalman tracking (Figure S7). Quantitative comparison of tracking and position estimation errors indicate that for the majority of cases COBWEBS tracking performance matches or exceeds that of Kalman tracking (Figure 4). Statistically insignificant data (p>0.05 for t test with Bonferroni multiple comparison correction) and cases where no trajectory was successful for either algorithm are not displayed. In regions where the plot is purple (right of dashed contour), Kalman tracking results in lower error than COBWEBS tracking. These regions have signal only OCRs of ~931 kcps or higher which is outside the fluorescence intensity of most readily accessible fluorescent labeling strategies. All other cases, COBWEBS tracking results in lower or similar error. Increasing the diffusion coefficient from 2 to 6 μm^2^/s shows similar trends in relative performance however, the range where Kalman tracking results in lower error than COBWEBS tracking expands to include particles with high backgrounds and signal OCRs of ~406 kcps or higher (Figure S8). Additionally, the magnitude of improvement for particles with lower OCRs in high backgrounds, the region where COBWEBS tracking is stable but Kalman tracking is not, is larger for D = 6 μm^2^/s than for the D = 2 μm^2^/s particle.

**Figure 4.**
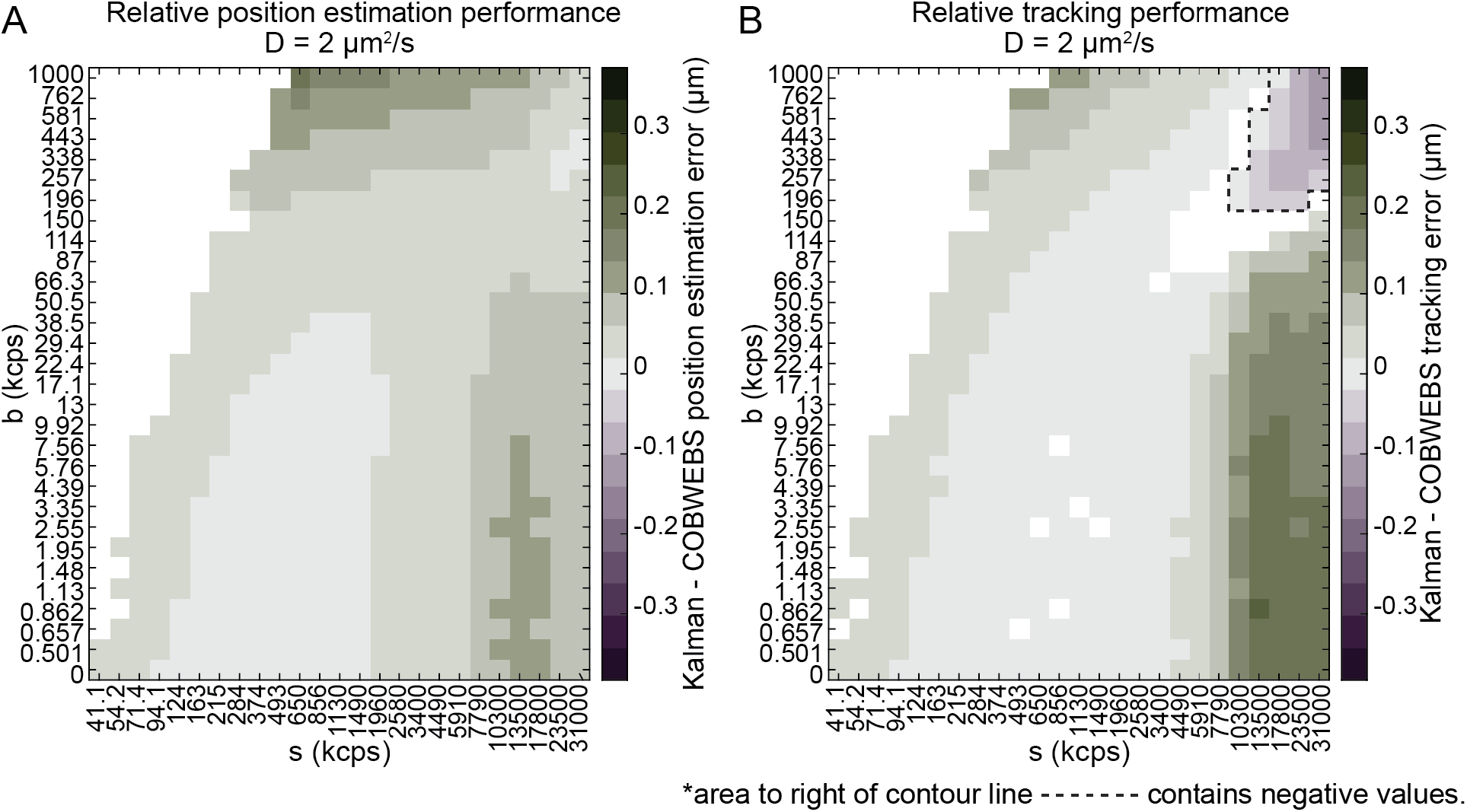
Relative error of Kalman and COBWEBS tracking over range of signal and background values. Positive values indicate lower error from COBWEBS tracking (green) while negative values enclosed by dashed contour line indicate lower error from Kalman tracking (purple). Statistically insignificant data (p>0.05 for t test with Bonferroni multiple comparison correction) and cases where no trajectory was successful for either algorithm are not displayed. (Population of mean error value for 10 trials, maximum duration 1 second)

An additional complication exists for particles with an exceptionally high OCR. At higher diffusivity, D = 6 μm^2^/s, tracking begins to fail at high signal values (OCR of ≥2000 kcps) for COBWEBS tracking (Figure S6). This corresponds to signal and background estimation performance decline not failure of Bayesian localization. Providing COBWEBS tracking algorithms with perfect information on the true *s* and *b* values restores functional tracking at high signal values (Figure S6). This breakdown was not observed in D = 2 μm^2^/s trials.

### C. Particle tracking performance in non-uniform background environments

Homogenous backgrounds are only part of the story. If one imagines a seesaw and the background photons as weights placed on the ends of the seesaw, it is intuitive to understand why a significant background perched on only one end of the seesaw becomes problematic. Since the Kalman estimates are Gaussian by definition, skewed backgrounds introduce bias which cannot be incorporated into position estimation. In contrast, the Bayesian particle probability distribution used in COBWEBS has no requirement for symmetry and so can more accurately model possible particle positions. In summary, uneven background intensities are more perturbative to Kalman estimation than to COBWEBS. This becomes experimentally relevant for a virus interacting with the complex environment of a cell (Figure 5A).

**Figure 5.**
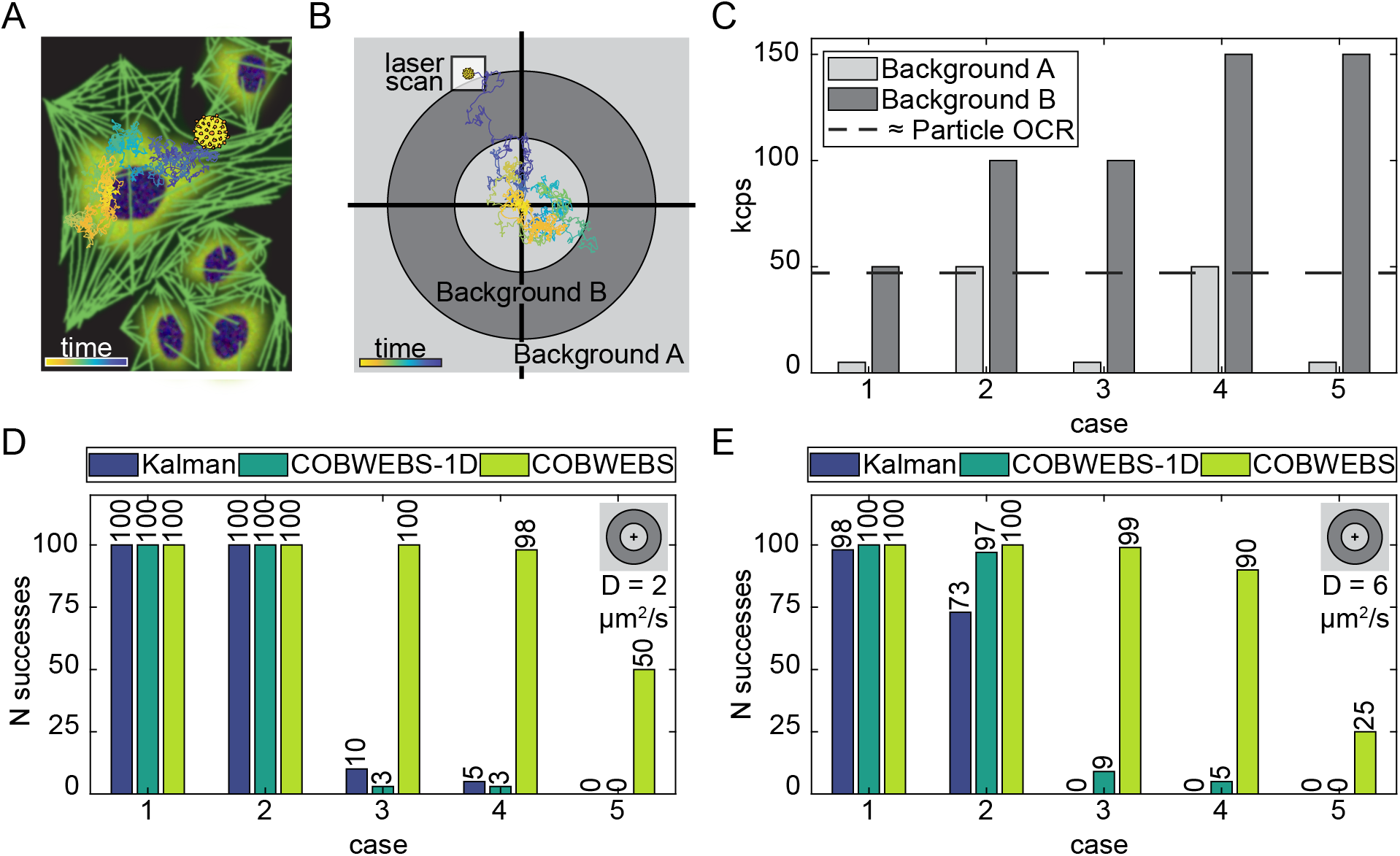
Uneven Background Simulations. **A.** Cartoon of a diffusive trajectory of a virus overlapped onto a cartoon of a fluorescently labeled cell. **B.** Cartoon of the donut structured background containing two distinct background interfaces (located at 2 and 4 μm from the initial particle position). The white box is the size of the laser scan size, drawn to scale. **C.** High and low background levels used for tracking simulations. Color of bar corresponds to the location of each background level as depicted in B. For example, in case 1 the background starts at 5k then increases to 50k. Cases are organized first by increasing jump size and second by initial background level. **D.** Donut success counts. (100 trials, 10 second max duration, *s* = 516.64 kcps, D = 2 μm^2^/s) **E.** Fast donut success counts. (100 trials, 10 second max duration, *s* = 516.64 kcps, D = 6 μm^2^/s)

To introduce the complexities of a nonuniform background environment to simulations, a radially symmetric donut background pattern was implemented (Figure 5B). The donut pattern enabled simulation of continued interactions with high and low interfaces in background level over a 10 second diffusive trajectory. A simplified dot pattern containing only a single interface at 1.5 μm was used to explore additional cases (SI Note 5.2). Simulations using patterned backgrounds suggest that the COBWEBS tracking approach increases trajectory durations in unevenly structured background environments.

First, a simulated particle with an OCR of 47 kcps and a diffusion coefficient of 2 μm^2^/s, consistent with a slow virus like particle (VLP) was explored. Even background simulations confirmed that all three algorithms could track particles at all background intensities of interest (Figure S10B). In uneven background environments all three algorithms work well for slowly diffusing VLPs provided the jump in background intensity is similar to the approximated OCR (Figure 5D, Case 1-2). The increasingly extreme examples of uneven background in Cases 3-5 show dramatically increased stability of COBWEBS position estimation (Figure 5D). The reproducibility of observed tracking success rates from Cases 3-5 (Table S4) show that the difference was significant.

In even background comparison simulations, increasing diffusive speed by a factor of three leads to a drop off in in Kalman performance at a background count rate at or above 100 kcps, but COBWEBS position estimation remains unaffected for this VLP (Figure S10C). For a faster particle, COBWEBS tracking displays significantly improved stability compared to Kalman for cases 1-4 (Figure 5E). Not only does COBWEBS estimation track particles beyond the Kalman performance drop off, but this effect extends to interfaces in background intensity up to at least ~3x the observed particle intensity.

The tracking benefit of COBWEBS is not dependent on initializing tracking in a high background environment. Although the raw success rate is higher for trajectories which begin in a high background environment and diffuse towards a lower background, independent of initial background intensity, COBWEBS trajectories are equally or more stable than their Kalman counterparts (Figure S10E, F). All dot cases (Figure S10E, F) demonstrate equivalent or higher overall success rates than their corresponding donut case (Figure 5D) likely due both to the shorter trajectory max duration of only 2 seconds and the reduced number of background interfaces.

Large errors in position estimate correlate spatially with the change in background interface and temporally with trajectory failure (Figure S9). Interestingly, the largest errors occur on the low background side of the interface. This suggests that transitions from low background into high background are more problematic than their high to low counterparts consistent with the higher failure rates observed in low to high dots in FigureS10E.

The superiority of COBWEBS position estimation at background interfaces extends to several particle intensities (Figure S11). The brighter case (OCR 93 kcps) can reliably track with any algorithm provided the difference in intensity at the interface is small (~1x particle brightness, Figure S11D). However, as the background intensity gap increases (cases 3-5), COBWEBS tracking again becomes significantly more stable than Kalman (Figure S11D). In homogenous background environments, COBWEBS tracking of dim particles occasionally fails at background levels as low as the approximate OCR of the particle (Figure S11A). Despite these occasional failures in the even background environment, introduction of a background interface shows superior COBWEBS tracking for cases (3-5) containing background levels and equivalent tracking for cases (1-2) (Figure S11C).

The computationally simplified COBWEBS-1D variant shows a stability benefit only for the fast VLP (Figure 5E) and the bright particle (Figure S11D) and in fact appears less stable than the Kalman tracking option for all other particles explored.

### D. Impact of diffusion coefficient parameter estimate mismatch on tracking performance

All three algorithms include an assumed diffusion coefficient. The impact of mismatches between this assumed parameter and the actual value are explored for the even background simulations with the faster (D = 6 μm^2^/s) VLPs. For the VLP, COBWEBS tracking works over a higher background intensity range than the Kalman tracking given perfect information on the diffusion coefficient (Figure 6). Kalman tracking duration declines for above 100 kcps background (~2x OCR), while COBWEBS tracking does not begin to decline until the background level reaches 200 kcps (~4x OCR) for the VLP (Figure 6). At non-zero background Kalman tracking benefits slightly from underestimation (dashed lines) of the diffusion coefficient as previously reported by Fields and Cohen (Figure 6A).^22^ In contrast, COBWEBS tracking is significantly impaired by underestimation of the true diffusion coefficient (Figure 6B). Contrastingly, overestimation of the diffusion coefficient by an order of magnitude for the VLP (dotted lines) decreased Kalman trajectory durations, while COBWEBS performance is unaffected. This indicates that the severe consequences of underestimation can be experimentally avoided by routinely overestimating the diffusion coefficient or simply setting the estimated diffusion coefficient to the upper trackable speed limit of a given tracking system.

**Figure 6.**
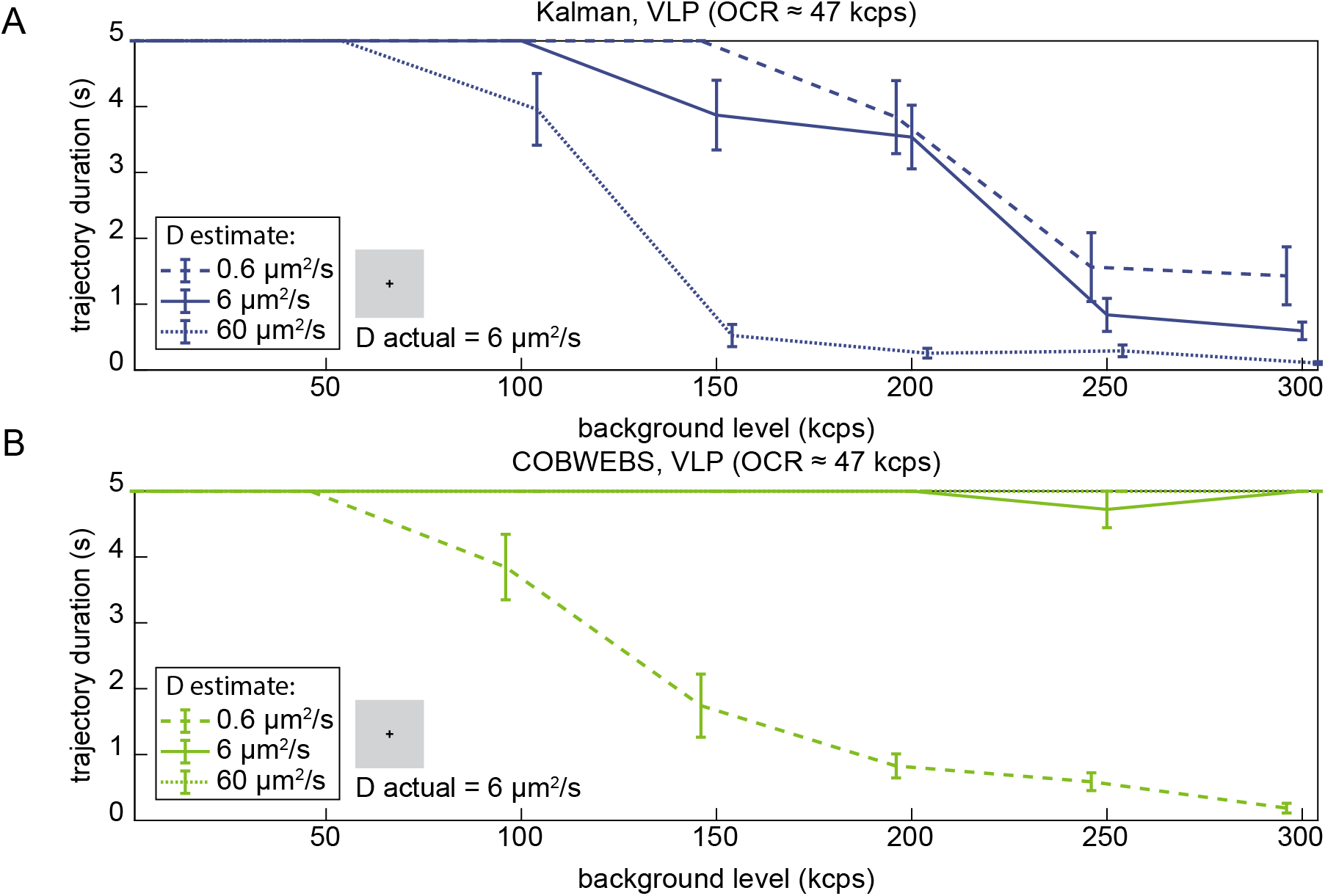
Diffusion parameter mismatch impact on trajectory duration for virus like particles. Diffusion coefficient estimates fed into localization were overestimated and underestimated by an order of magnitude for **A.** Kalman and **B.** COBWEBS tracking (10 trials, 5 second max duration, *s* = 516.64 kcps, D actual = 6 μm^2^/s). Error bars are standard error of the mean.

These trends are maintained for additional particle intensities. For a bright particle, in COBWEBS tracking, overestimation of the diffusion coefficient does not affect trajectory duration at all (Figure S12B). Further, even with reduced COBWEBS performance due to underestimation of the diffusion coefficient it still outperforms Kalman’s best case scenario (Figure S12B). For the dim particle, COBWEBS tracking works over a more limited range of background values than Kalman estimation (Figure S12A). For the dim particle overestimation appears to improve trajectory durations. Increasing the diffusion coefficient estimate artificially increases uncertainty in particle positions suggesting that for dim particles especially in high background environments, COBWEBS tracking becomes too “confident” about particle locations. This could be due to simply not having enough available photons with which to make good particle position estimates. For limited diffusive speeds, impaired tracking at low count rates can be restored by extending the bin time (SI Note 7). For the dim particle, underestimation reduces trajectory duration consistent with the trends from the other two intensities.

## IV. Discussion

COBWEBS position estimation dramatically improves tracking robustness to inhomogeneous backgrounds in simulated trajectories. COBWEBS position estimation reduces position estimation error and tracking error for the majority of particle and background intensity combinations. Tracking error is important both for particle retention and for data interpretation since stage positions are routinely used as particle positions due to computational complexity of post processing data. COBWEBS tracking potentially extends the range of background levels in which tracking is possible for particles of adequate brightness. Additionally, the delineation between trackable and non-trackable particles is more invariant with diffusive speed than that observed in Kalman position estimation indicative of improved consistency. Most importantly, COBWEBS tracking dramatically increases possible trajectory duration for freely diffusing particles in uneven background environments corresponding to the environment of a live cell. Finally, COBWEBS position estimation is more resilient to overestimation of diffusion coefficients than Kalman position estimation. Provided that they could develop a suitable signal and background estimation strategy or set reasonable estimates for signal and background levels a priori, this adaptive algorithm could be extended to work in other active feedback methods such as orbital tracking and tetrahedral tracking, therefore expanding generally the ability to track diffusive particles in complex environments.

Before the experimental benefits of COBWEBS tracking can be realized it must be translated onto an FPGA, which has limited computational resources for complex mathematical functions such as multiplication and division. By converting all probability distributions to logarithmic space, almost all multiplications and divisions can be converted into addition and subtraction reducing the computational complexity of COBWEBS estimation. The largest obstacle then becomes the numerical convolution accounting for diffusion, which must be performed in real space. The conversion between logarithmic and real space can be performed using look up tables, as can the factorial from Eq. 6. The exponential term of *γ_k_* (Eq. 5) depends only on laser scan positions and particle position. This partial *γ_k_* can also be stored as a look up table for each possible laser position reducing calculation of the expected count rate for a given position pixel to a single multiplication by *s_Est_*(*k*) and addition of *b_Est_*(*k*).These simplifications reduce COBWEBS estimation to contain minimally complex operations. Further reducing the grid sampling would reduce both the overall number of elements computed per estimate and the size of the diffusion kernel. Therefore, given the use of allowable operations and customizability of estimation grid inter-point spacing, it is reasonable to expect that given an adequate FPGA it will be possible to implement this two-dimensional COBWEBS in real time.

The same cannot be said for a three-dimensional variant of COBWEBS which simply introduces too many operations to fit into available computational resources. Expanding the probability grid to the third dimension would increases the number of elements which must be stored and updated per estimate from *V*^2^ to 2 · *V*^3^ (since the Z axis scan width is 2x the size of the X, Y knights tour). Additionally, each of those 2 · *V*^3^ points would require 3 · *kernel size* multiplications per point updated compared with the 2 · *kernel size* multiplications of COBWEBS. Should FPGA technology improve dramatically, the true 3D variant may become experimentally possible. In the meantime, COBWEBS can be combined with traditional estimation approaches along the Z axis and provide a stability benefit in uneven background environments even if that stability is only along two of the three tracked axes.

Overall, the COBWEBS algorithm is robust to large and abruptly changing signal-to-background ratios, making it a prime candidate for advancing active-feedback single-fluorophore tracking to the complex cellular interior as well as other challenging tracking environments.

