## Supplementary Methods and Data for "Combined online Bayesian and windowed estimation of background and signal localization facilitates active-feedback particle tracking in complex environments"

#### SI Note 1: Simulation Details

##### 1.1 Particle motion

The mean value of displacement for random diffusion along a single axis is approximated by a normal distribution with a mean displacement of 0 and a variance of $2D\tau$ where $D$ is the diffusion coefficient and $\tau$ remains the bin time.[1]

$$\Pr(x) = a * \exp\left(\frac{-|x]^2}{4D\tau}\right) \quad \text{Eq. S1}$$

Here, $a$ is a normalization/prefactor which we have not analytically defined due to eventual ongoing numerical normalization.

Particle trajectories were generated using pseudo-random numbers from a normal distribution with variance $2D\tau$ (SI Eq. S1). The cumulative sum of all pseudo-random steps taken gives the particle's position at a given timepoint. Bin time ($\tau$) was 20 μs (unless otherwise specified) and experimentally relevant diffusion coefficients of either 2 or 6 μm$^2$/s were selected.

##### 1.2 Stage control and response

Experimentally, 3D-SMART utilizes an integral feedback controller where the position estimates are integrated, multiplied by a constant, and fed into the stage control loop. For this work, we used a more general PI controller as described below:

$$Command_f = k_p \cdot x_f + k_i \sum_{n=1}^{f} x_f \quad \text{Eq. S2}$$

The stage command for a given bin f is given by Eq. S2 where $x_f$ is the position estimate for a given bin and $k_i$ and $k_p$ are integral and proportional control constants which were numerically optimized. Stage positions resulting from a given command were simulated by convolving the stage command sequence output from Eq. S2 with the empirically determined step response function as previously described.[2] Convolution computation duration for stage simulation was improved using Fourier convolution.

Experimentally, 3D-SMART has previously used integral-only control.[2, 3] Integration accounts for Kalman position estimate fluctuations around the particle's true position. Since COBWEBS position estimates fluctuate less, it was possible that proportional control, where the estimate for each given bin is used as a component of the stage command sequence would be useful.

Optimizing $k_i$ and $k_p$ values for each particle would be temporally prohibitive in both simulations and experiments. Therefore, "globally optimized" feedback parameters were generated using selected cases. Optimal values for all cases included are in Table S1. An example optimization is shown in Figure S2 to show the high variability in acceptable $k_i$ and $k_p$ parameters. The wide range of acceptable values for individual cases reinforces the ability to choose a single globally optimized feedback parameter.

Using globally optimized feedback parameters, in COBWEBS tracking the proportional control term accounts for less than 10% of the overall control signal at least 90% of the time (Figure S1). COBWEBS tracking control remains dominated by the integral term despite increased accuracy of the COBWEBS position estimation. Kalman simulations showed no benefit to inclusion of the proportional control term and so Kalman tracking was performed using integral control alone.

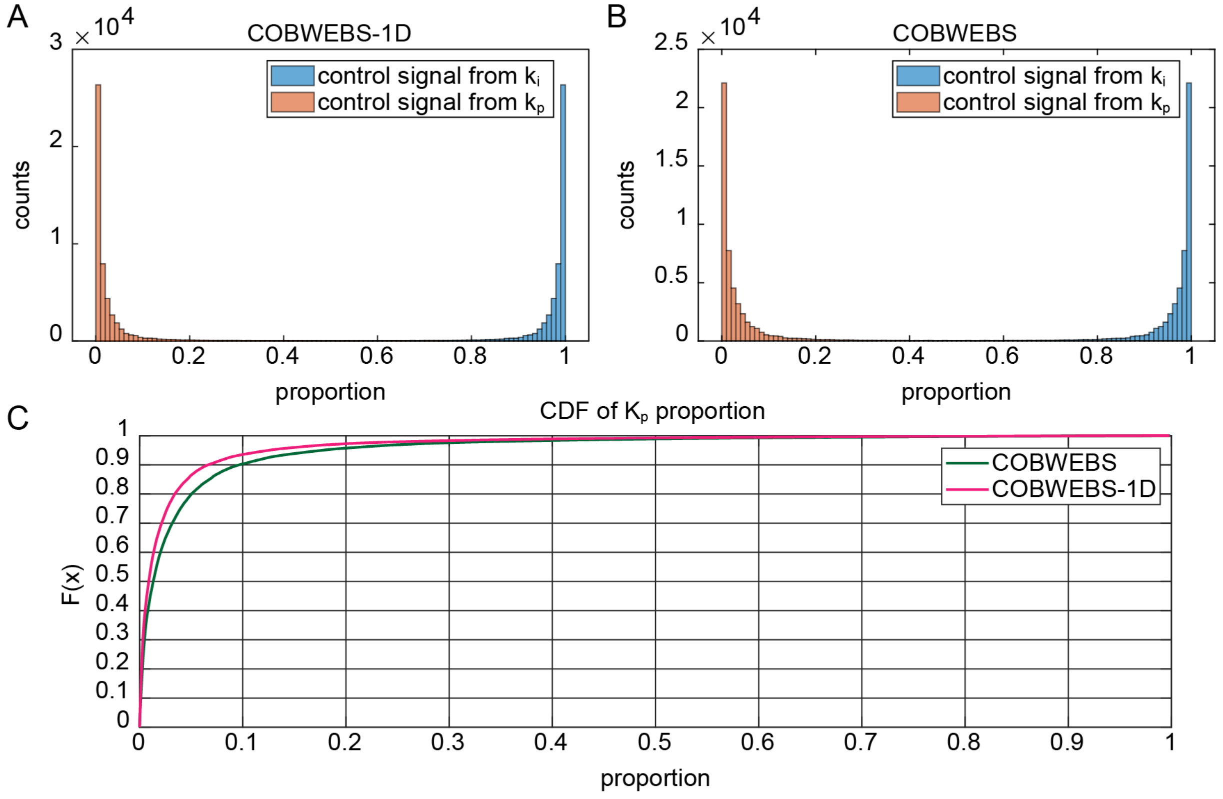

**Figure S1. Relative contribution of $k_i$ and $k_p$ to overall control signal. A, B.** Histogram of proportion of control signal originating from $k_i/k_p$ along x axis in example trajectory 1/25 (s = 516.64 kcps, b = 50 kcps, D = 2 μm$^2$/s, duration = 1 s) from Figure 3 for both 1D COBWEBS (A) and 2D COBWEBS (B) position estimation. **B.** Empirical CDF version of $k_p$ proportion histogram from A, B.

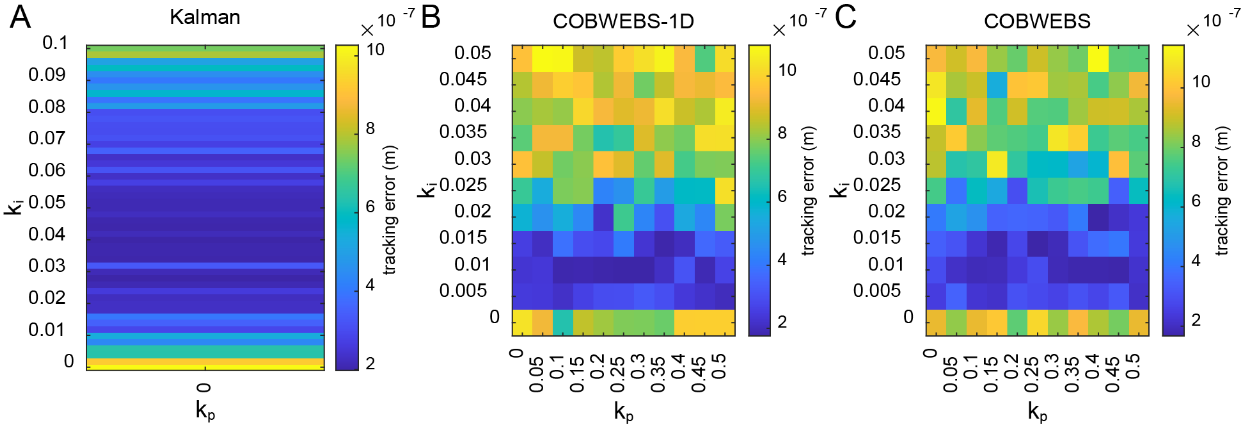

**Figure S2*. Feedback parameter optimization.* A.** Mean Kalman tracking error for given $k_i$. **B.** Mean 1D-COBWEBS tracking error for given combination of $k_i$ and $k_p$. **C.** Mean 2D-COBWEBS tracking error for given combination of $k_i$ and $k_p$. (s =1 23.969 kcps, b = 29.367 kcps, trials per $k_i$ = 10, duration = 0.1 sec). The feedback parameter at the global minimum tracking error value was deemed the optimal value for a given case.

**Table S1. Optimal feedback parameter values.** Feedback parameter ($k_i$, $k_p$) values with minimum tracking error over 10 trials 0.1 sec in duration per $k_i$ $k_p$ combination tested.

| D ($\mu m^2$/s) | s (kcps) | b (kcps) | s:b ratio | Kalman $k_i$ | COBWEBS-1D $k_i$ | COBWEBS-1D $k_p$ | COBWEBS $k_i$ | COBWEBS $k_p$ |
|---|---|---|---|---|---|---|---|---|
| 1.00E-12 | 493 | 51 | 10 | 0.024 | 0.015 | 0.25 | 0.020 | 0.40 |
| | 10273 | 1000 | 10 | 0.038 | 0.015 | 0.05 | 0.025 | 0.15 |
| | 1129 | 10 | 114 | 0.020 | 0.020 | 0.20 | 0.020 | 0.45 |
| | 124 | 0 | Inf | 0.018 | 0.020 | 0.35 | 0.015 | 0.50 |
| | 10273 | 0 | Inf | 0.018 | 0.015 | 0.05 | 0.025 | 0.15 |
| 6.00E-12 | 493 | 51 | 10 | 0.040 | 0.020 | 0.10 | 0.025 | 0.15 |
| | 10273 | 1000 | 10 | 0.074 | 0.020 | 0.00 | 0.025 | 0.00 |
| | 1129 | 10 | 114 | 0.028 | 0.025 | 0.00 | 0.025 | 0.10 |
| | 124 | 0 | Inf | 0.026 | 0.020 | 0.20 | 0.025 | 0.25 |
| | 10273 | 0 | Inf | 0.018 | 0.015 | 0.05 | 0.020 | 0.00 |
| | 124 | 29 | 4 | 0.040 | 0.010 | 0.30 | 0.020 | 0.40 |
| | 18 | 1 | 36 | 0.008 | 0.005 | 0.40 | 0.005 | 0.45 |
| | 30997 | 0 | Inf | 0.018 | 0.020 | 0.00 | 0.020 | 0.00 |
| 2.00E-11 | 493 | 51 | 10 | 0.048 | 0.020 | 0.10 | 0.025 | 0.10 |
| | 10273 | 1000 | 10 | 0.100 | 0.020 | 0.00 | 0.025 | 0.00 |
| | 1129 | 10 | 114 | 0.030 | 0.020 | 0.05 | 0.025 | 0.05 |
| | 124 | 0 | Inf | 0.026 | 0.025 | 0.15 | 0.025 | 0.15 |
| | 10273 | 0 | Inf | 0.020 | 0.020 | 0.00 | 0.020 | 0.05 |
| Mean | | | | 0.0330 | 0.0181 | 0.125 | 0.0217 | 0.186 |
| Standard Deviation | | | | 0.02 | 0.005 | 0.1 | 0.005 | 0.2 |
| Optimized Value | | | | 0.032 | 0.02 | 0.1 | 0.02 | 0.15 |

##### 1.3 Simulated photon generation

During a simulated trajectory, for each bin, the expected rate of photon generation for the input s and b parameters was calculated using Eq. 5 and the displacement between the particle and laser spot centers. Then, for each bin, pseudo-random numbers from the calculated Poisson distribution from Eq. 6 became the number of photons detected in a given bin. The point spread function sigma was modeled with a width of 150 nm. For explorations over a range of values, s varied logarithmically from 41.1 to 30997 kcps and b values varied logarithmically from 0 to 1000 kcps.

For theoretical development of signal and background estimation, the displacement between the particle and laser spot centers were modeled as pseudo random numbers from a normal distribution (σ = 0.1 μm, μ = 0).

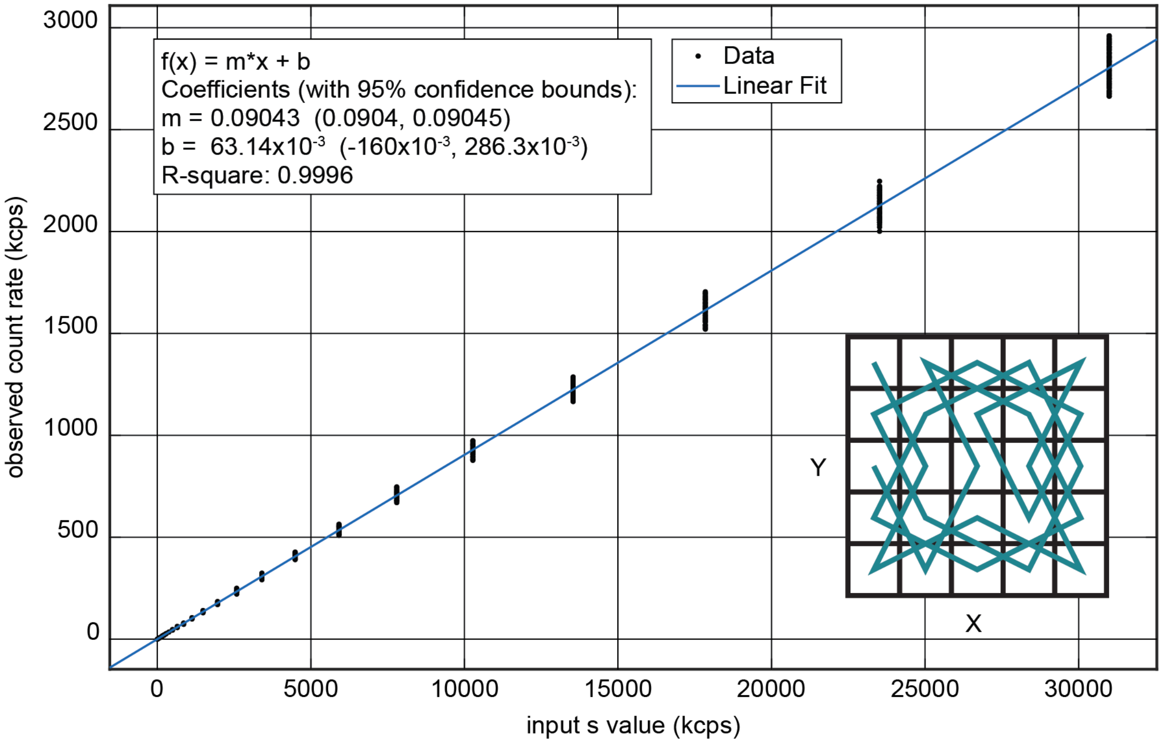

**Figure S3. Linear fitting of OCR to input s value.** Data from same b=0 photon data as Figure 2A but with OCR averaged over entire laser knight's tour scan pattern (Inset) instead of a single pixel type. (b = 0, 500 trials, 5000 bins (100 ms) of data per trial, Particle positions = Normal (σ = 0.1 μm, μ = 0)).

During tracking, observed photon numbers are reduced compared to the theoretical max count rate s. In practice the actual OCR will vary with tracking performance due to changes in the offset between particle position and scan center. However, approximating displacements as a normal distribution generating a uniform conversion to predict the observed count rate for a given s value enables a priori selection of experimentally relevant particles. Since photons are generated without assignment to a source, all OCRs discussed in this paper were computed by multiplying the input s value by 0.09043.

#### 1.4 Conversion from analytical Bayesian to numerical Bayesian

For real-time tracking a numerically efficient implementation is needed. In the present work, particle positions are gridded into 5 nm. This spacing is smaller than the expected diffusive step size $\sqrt{2D\tau}$, which for the fastest particle studied here is 15.5 nm. Additionally, the particle's position is assumed to lie within the overall scan area of the laser (1 µm x 1 µm) thus limiting the overall grid of potential particle positions to a computationally feasible 201 × 201 grid.

The mathematically complete Bayes theorem requires scaling the probability by the probability of detecting n photons over all space (denominator in Eq. 7). Here we simply assume the particle is within the scan area, the total probability must equal one and calculate and apply the appropriate numerical normalization constant in a second computational step rather than calculating the analytical normalization constant.

Computational speed for calculation of the rate of expected photon detection was improved by using a look-up table (LUT) for the signal and background estimate invariant portion of $\gamma_k$. The exponential term of $\gamma_k$ (Eq. 5) depends only on laser scan positions and particle position. Laser scan positions are restricted to a 5 × 5 grid. Therefore, a LUT containing the 201 × 201 grid of partial $\gamma_k$ values was generated for each of the 25 possible laser positions.

#### 1.5 Numerical convolution for diffusion and simplifications

The two-dimensional convolution with particle diffusion is one of the more computationally intensive steps in position estimation. Two simplifications are possible.

First, the symmetry of the diffusion kernel leads to the ability to separate the diffusion convolution kernel into two sequential convolutions along the x and y axes respectively reducing the total number of necessary multiplications per pixel from $V^2$to $2V$where $V$ is the size of the diffusion kernel in pixels. Using the separated diffusion kernel the update equation becomes the numerical convolution between m and columns then m and the rows of $f_{k-1|k-1}$ where q runs over all subscripts that lead to legal matrix indices.

$$g_{k-1|k-1}(x,y) = \sum_q m(q) f_{k-1|k-1}(:,y-q) \qquad \textit{Eq. S3}$$

$$f_{k|k-1}(x,y) = \sum_q m(q) g_{k-1|k-1}(x-q,:) \qquad \textit{Eq. S4}$$

To prevent drift in the size of the probability matrix grid, the position probability matrix is zero padded around the edges prior to convolution and the output is clipped to return a matrix of the same size as the input. The technical invalidity of the convolution around the edges of the particle position estimation grid reduces the likelihood of predicting a particle position at the edge of the area. However, in a well tracked particle the particle spends minimal time near the edges of the scan area minimizing the relevance of this invalidity.

Second, while the diffusion kernel is defined over all possible values of x and y, in practice its size can be limited to a much smaller number of non-zero terms. For numerical simplicity, only values of the diffusion kernel above 10% of the max value were used. This results in tractable diffusion kernel sizes of 7 and 13 pixels for particles with D = 2 and 6 µm$^2$/s respectively. These diffusion kernels contain 97% of the total step size probability distribution. The clipped kernel produces convolutions similar to ones performed using the complete diffusion kernel, with the benefit of significantly decreased computation time.

#### SI Note 2: One-dimensional variant, COBWEBS-1D

COBWEBS-1D estimation uses a simplification of the 2D Bayesian estimation method derived above which assumes that the probability distribution of particle positions along axes are entirely independent. This converts the numerically discretized priors and posteriors into two independent vectors of length 201.

No diffusion kernel clipping was required to make COBWEBS-1D position estimation computationally feasible, so the entire kernel was used. As such the numerical convolution to calculate is performed along single axes instead of all columns followed by all rows sequentially.

$$f_{k|k-1}(x) = \sum_{q} m(q) g_{k-1|k-1}(x-q) \quad \textit{Eq. S5}$$

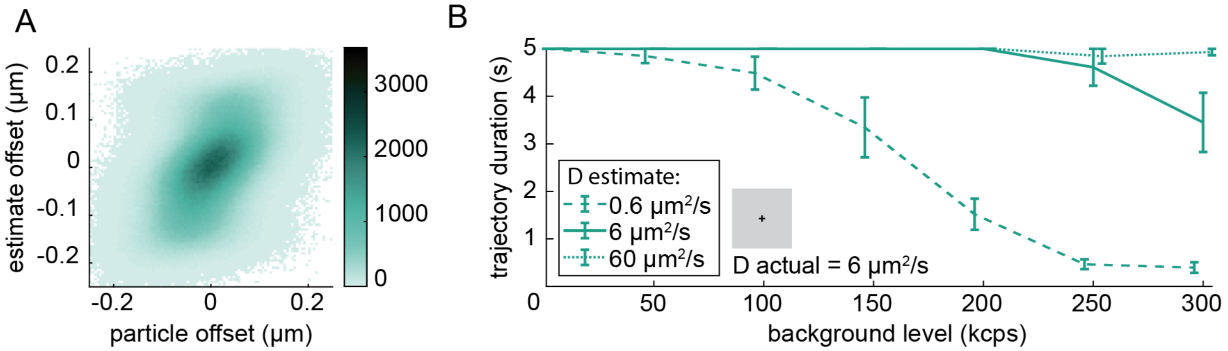

**Figure S4. COBWEBS-1D Performance. A.** COBWEBS-1D equivalent panel from Figure 3D. Correlation coefficient between actual offset and estimated position is 0.4873 for COBWEBS-1D. **B.** COBWEBS-1D equivalent panel to Figure 6. Same experimental conditions apply.

COBWEBS-1D position estimation results in a slightly worse overall position estimation error than the Kalman position estimation (Figure 3C). However, somewhat paradoxically despite a slightly higher estimate error, the tracking error of COBWEBS-1D estimation is lower than Kalman estimation. This discrepancy is explained by our "responsivity" metric of correlation coefficients (Figure S4A). Moving from Kalman estimation to COBWEBS-1D increased the correlation coefficient between actual and estimated offset from 0.45 to 0.49. The slightly improved strength of linear relationship between estimated and actual particle positions explains why the COBWEBS-1D variant demonstrates an improved tracking error despite slightly reduced position estimation accuracy than Kalman. Of course, the COBWEBS correlation coefficient of 0.63 is much higher than both other options.

The COBWEBS-1D variant only preserves probability likelihood only along a single axis reducing the number of elements and overall 2D resolution of the prior. This reduction in retained information leads to the visually observable non-linear error components in the upper left and lower right-hand corners of the offset estimates where particles are estimated to be on the wrong side of the stage. These intermittent errors account for the increased position estimation error when compared to the Kalman. Overall, COBWEBS-1D estimation appears to be slightly more sensitive to particle motions than Kalman but also less accurate.

COBWEBS-1D tracking shows identical trends to COBWEBS tracking with respect to sensitivity to mismatches in estimated and actual diffusion coefficient parameters (Figure S4B). Underestimation of diffusion coefficient is catastrophic, but the algorithm displays improved resilience to significant overestimations in diffusion coefficient.

In general, despite the significantly reduced computational complexity of the COBWEBS-1D variant its assumption of independence in particle position likelihoods appears to cripple the approach leading to under-performance. In many cases, COBWEBS-1D underperforms the Kalman filter.

#### SI Note 3: Signal and Background Estimation

##### 3.1 Alternate combinations of pixels for estimation of signal and background

Six equivalent scan positions based on symmetry and distance from center of laser scan (Figure S5) were labeled as pixels A-F. The weighting factors (Figure 2A, B) for each possible combination of 2 pixel types were used to calculate the percent error associated with signal and background estimation using all possible combinations of 2 pixel types. The central and corner pixels led to the lowest overall estimate error. In general estimation strategies containing more central pixels were best at estimating signal values while more distal pixels are best for estimating background levels.

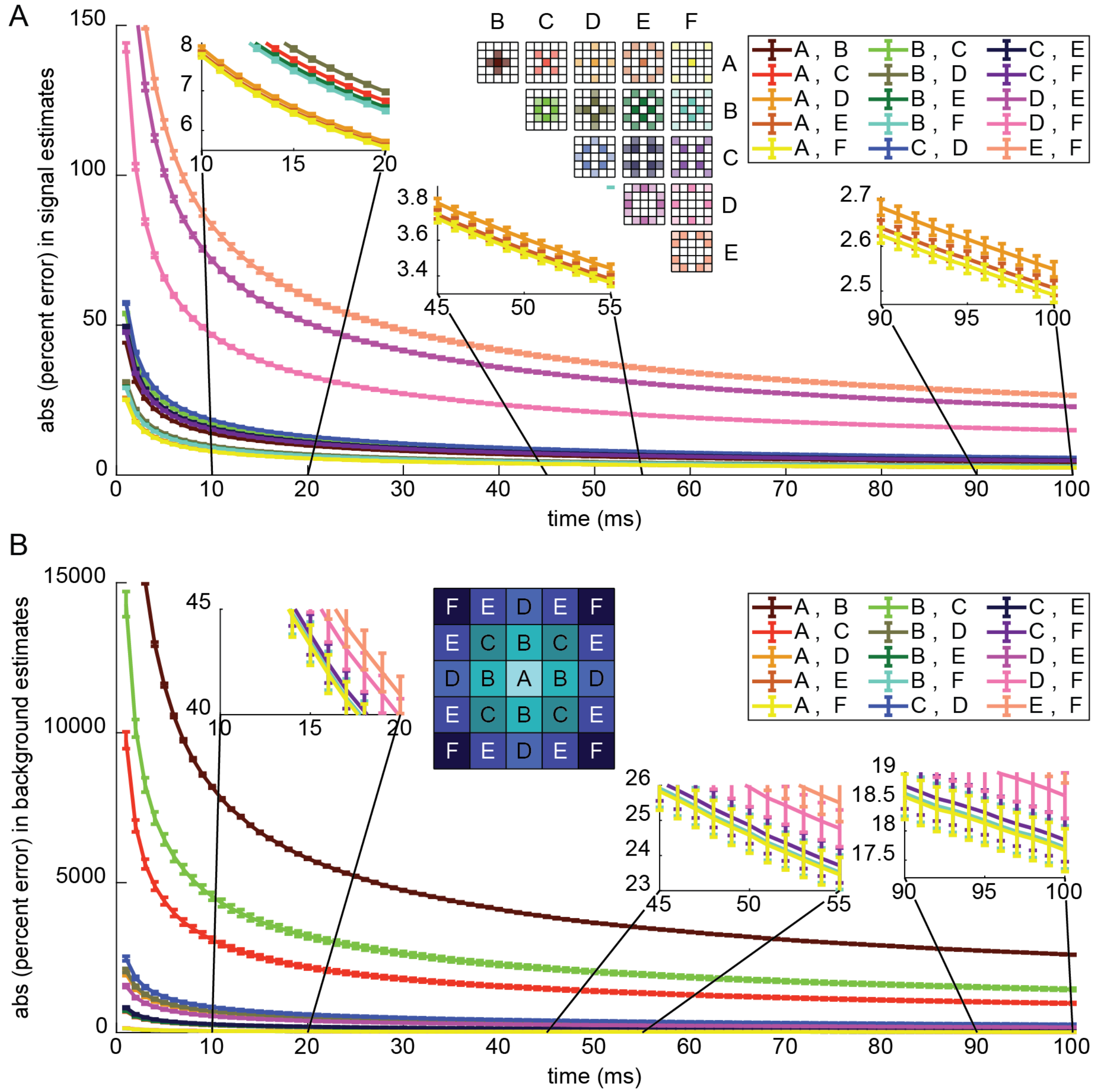

**Figure S5. Signal and background estimates for different pixel combinations versus time.** Overall absolute value of percent error for signal and background estimates. Error bars are standard error of the mean. Window sizes ranged from 1 to 100 ms (500 trials of 100 ms) with particle positions randomly generated using a normal distribution with standard deviation of 0.1 and mean of 0.

##### 3.2 Impact of windowed signal and background estimation on tracking in even backgrounds

A faster diffusing particle moves more quickly and thus is more difficult to hold in the center of a moving stage which responds at a fixed speed. It may be necessary to increase the scan area to improve contrast between signal and background contributions to improve ability to estimate the background level at higher diffusive speeds.

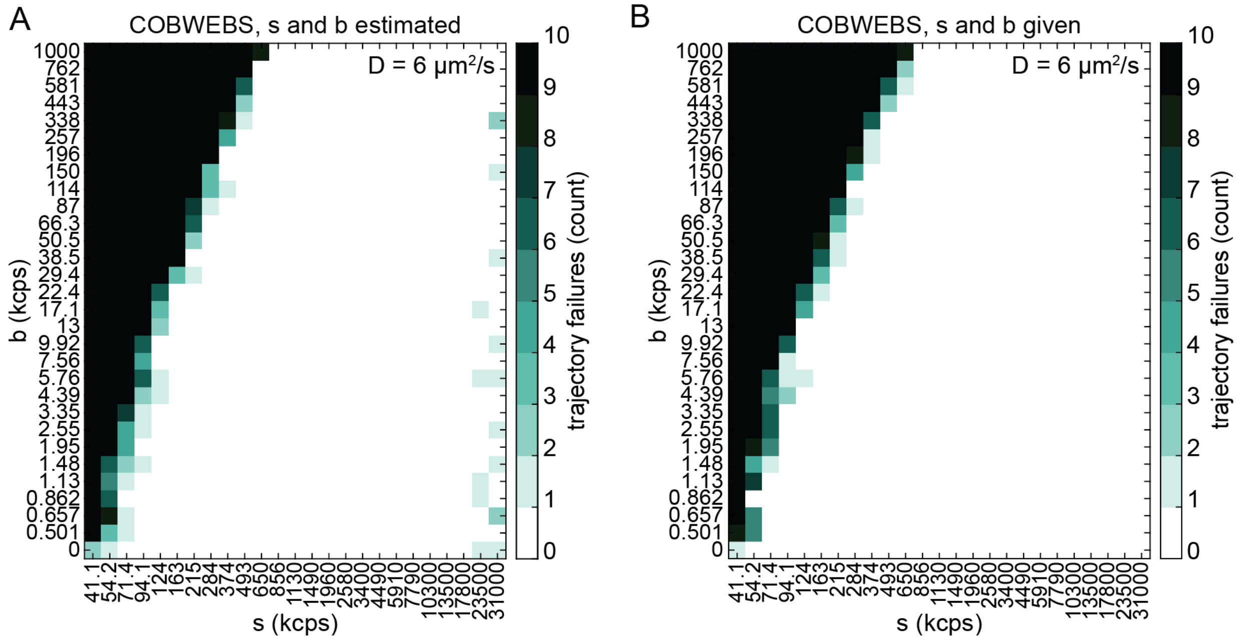

**Figure S6. Impact of signal and background estimation on tracking performance. A.** Number of failures out of 10 trials for each tested signal and background combination with the implemented signal and background estimation. (D = 6 μm$^2$/s, Duration = 1 sec). **B.** Number of failures out of 10 trials for each tested signal and background combination given perfect information about s and b values used to generate data. (D = 6 μm$^2$/s, Duration = 1 sec)

### SI Note 4: Particle tracking in even background environments

#### 4.1 Detailed statistics comparing all three algorithms from even background case.

**Table S2: ANOVA F values and calculated p value**

| ANOVA† | F(2, 74) | p |
|---|---|---|
| Position Estimation Error | 6580.95 | 3.04238e-82 |
| Tracking Error | 950.12 | 1.75968e-52 |

†Statistics generated using MATLAB function anova1

**Table S3: Tukey's t test and calculated p values for differences between algorithms**

| Tukey-Kramer† | Group A | Group B | Lower Limit | A-B | Upper Limit | P-Value |
|---|---|---|---|---|---|---|
| Position Estimation Errors | Kalman | COBWEBS-1D | -6.25E-09 | -5.55E-09 | -4.86E-09 | 9.56E-10 |
| | Kalman | COBWEBS | 2.50E-08 | 2.57E-08 | 2.64E-08 | 9.56E-10 |
| | COBWEBS-1D | COBWEBS | 3.06E-08 | 3.13E-08 | 3.20E-08 | 9.56E-10 |
| Tracking Errors | Kalman | COBWEBS-1D | 1.25E-08 | 1.35E-08 | 1.44E-08 | 9.56E-10 |
| | Kalman | COBWEBS | 1.52E-08 | 1.61E-08 | 1.71E-08 | 9.56E-10 |
| | COBWEBS-1D | COBWEBS | 1.72E-09 | 2.67E-09 | 3.62E-09 | 1.13E-08 |

†Statistics generated using MATLAB function multcompare.

##### 4.2 Position estimation error over range of particle brightness and background intensities for multiple diffusion coefficients

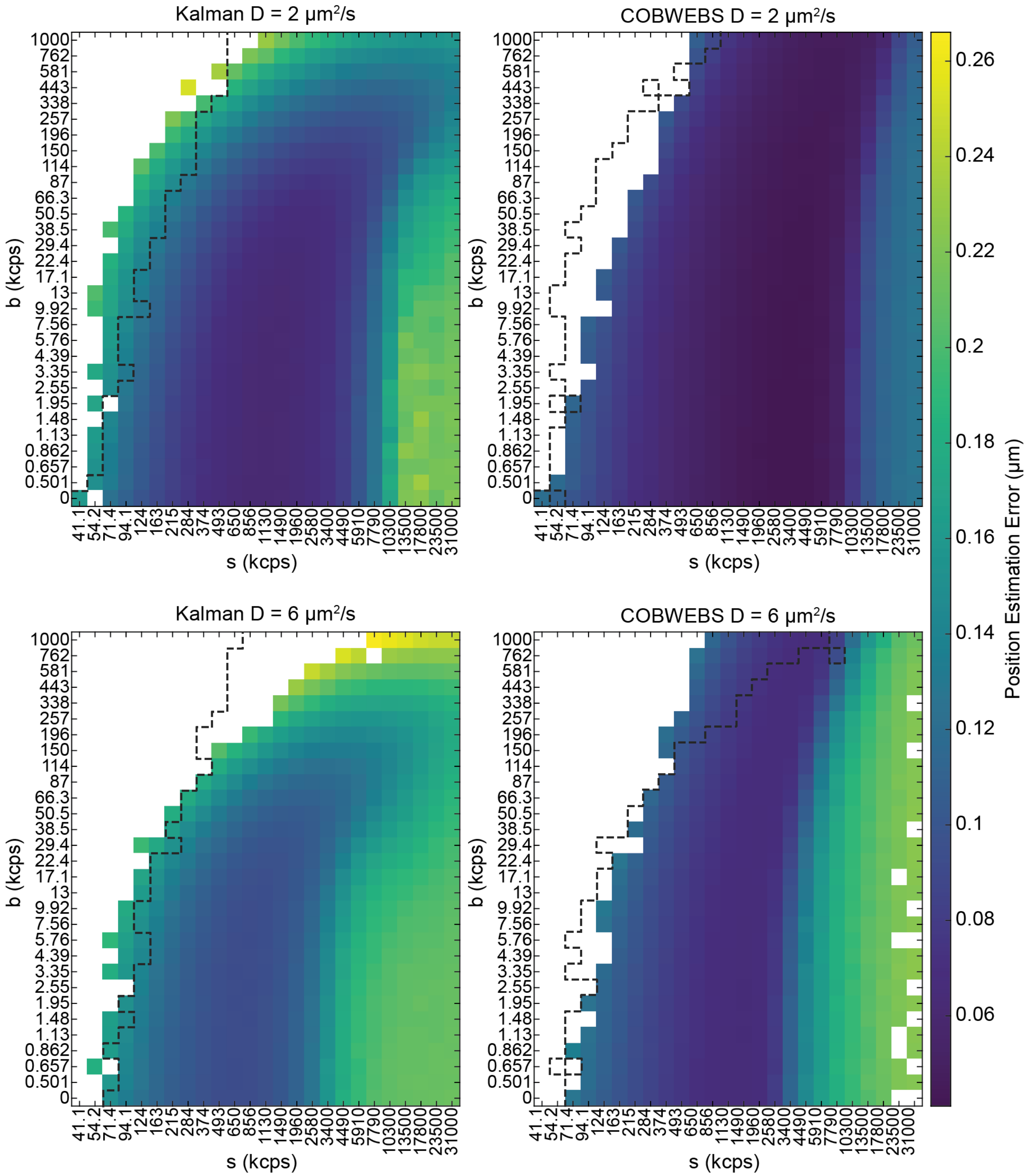

**Figure S7. Comparison of mean position estimation error between algorithms for a range of particle and background intensities.** Each signal and background combination contains 10 trials with a maximum duration of 1 second. Errors for trajectories where the duration of any trial was less than the full second are not displayed. Dashed lines in all plots are the edge of the trackable/not filter for the corresponding algorithm at the same diffusion coefficient.

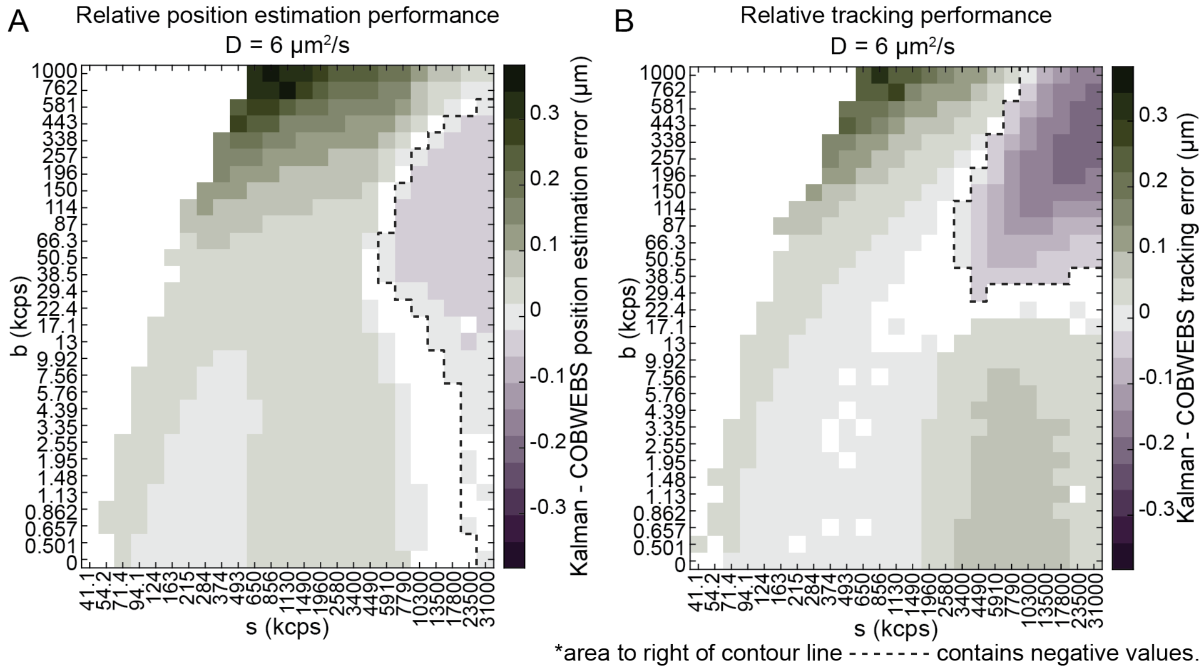

**Figure S8. Values of Kalman mean error minus COBWEBS mean error.** Positive values indicate lower error from COBWEBS tracking (green) while negative values enclosed by dashed contour line indicate lower error from Kalman tracking (purple). Statistically insignificant data ($p>0.05$ for t test with Bonferroni multiple comparison correction) and cases where no trajectory was successful for either algorithm are not displayed. (10 trials, maximum duration 1 second)

Tracking error is compared over all possible signal and background combinations filtered by whether tracking was reliably possible. For all particles with $s \geq 650$ kcps COBWEBS estimation tracks over a similar or extended range of background intensities for both diffusion coefficients (Figure S7). For $s < 650$ kcps there is a distinct cluster of cases where Kalman estimation appears to track in higher background intensities than COBWEBS (Figure S7). This cluster gets smaller when diffusivity increases 3-fold. The range of trackable particles in COBWEBS estimation is more consistent after increased diffusive speed than Kalman tracking; the boundary shifts less between the two COBWEBS cases than it does in the Kalman equivalent (Figure S7). COBWEBS tracking has a lower minimum error value than the corresponding Kalman cases indicated by the darker blue/error minimum value on the COBWEBS plots of Figure S7.

### SI Note 5: Particle tracking in inhomogeneous background environments

#### 5.1 Additional information about Figure 5

**Table S4: Reproducibility of tracking success in Figure 5D, VLP Cases 3-5.**

| $B_{low}$ (kcps) | 5 | | | 50 | | | 5 | | |
|---|---|---|---|---|---|---|---|---|---|
| $B_{high}$ (kcps) | 100 | | | 150 | | | 150 | | |
| | | COBWEBS | | | COBWEBS | | | COBWEBS | |
| | Kalm. | 1D | 2D | Kalm. | 1D | 2D | Kalm. | 1D | 2D |

| | | | | | | | | | |
|---|---|---|---|---|---|---|---|---|---|
| trial 1 (Figure 5) | 10 | 3 | 100 | 5 | 3 | 98 | 0 | 0 | 50 |
| trial 2 | 7 | 9 | 100 | 2 | 3 | 99 | 0 | 0 | 51 |
| trial 3 | 6 | 7 | 100 | 4 | 0 | 99 | 0 | 0 | 49 |
| Standard Deviation | 2.1 | 3.1 | 0.0 | 1.5 | 1.7 | 0.6 | 0.0 | 0.0 | 1.0 |

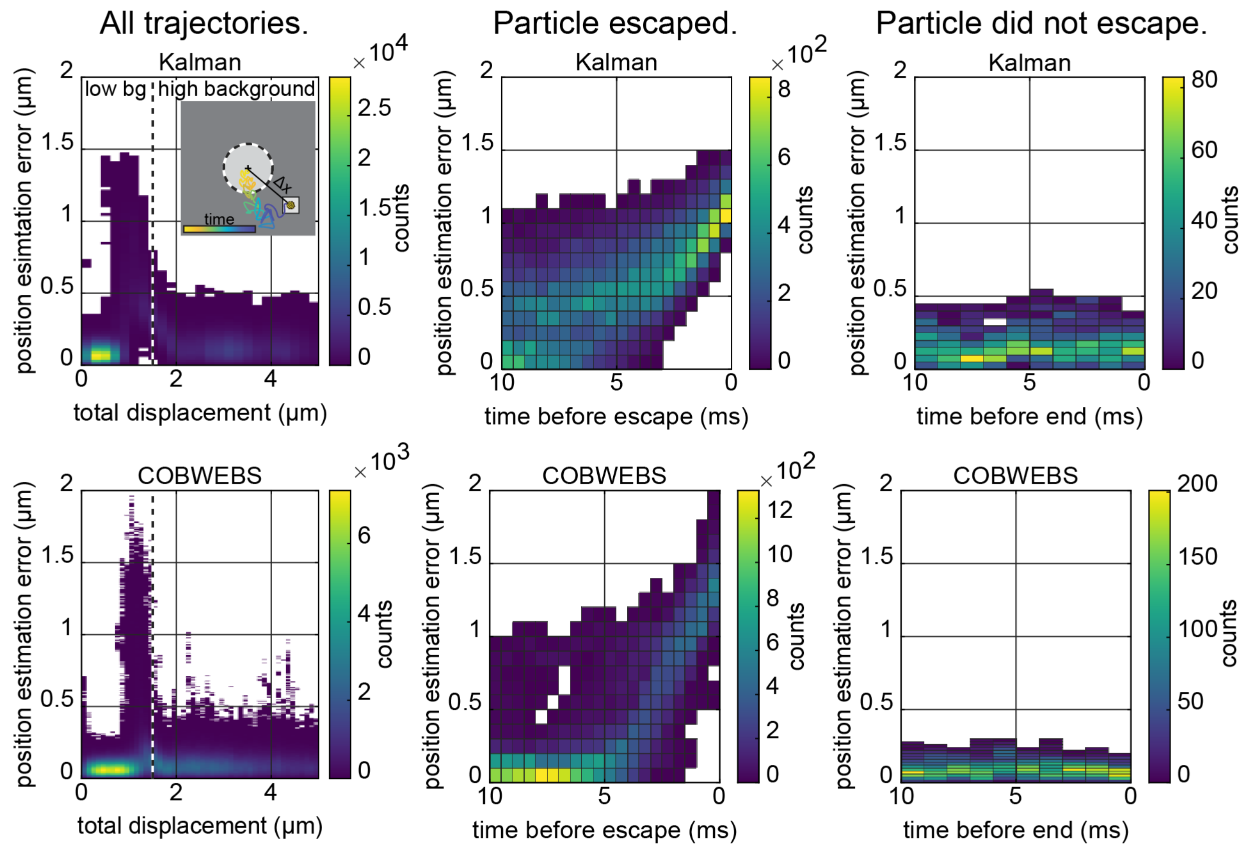

**Figure S9. Position estimation errors correspond to background interfaces and particle drops for all three algorithms.** (100 trials, max duration 2 seconds, s = 206.656 kcps, b low = 2 kcps, b high = 40 kcps, D = 2 μm$^2$/s). Row 1-Kalman. Row 2-COBWEBS. Column 1- position estimation error vs distance of particle from center of original position in a low to high dot with interface change at 1.5 μm (indicated by the vertical black and white dashed line). Column 2-position vs time before trajectory end in trajectories which ended prior to the maximum duration (failed trajectories). Column 3-position vs time before trajectory end in trajectories which lasted the full 10 seconds (successful trajectories).

##### 5.2 Even background comparison simulations, additional particle intensities, and additional background patterns

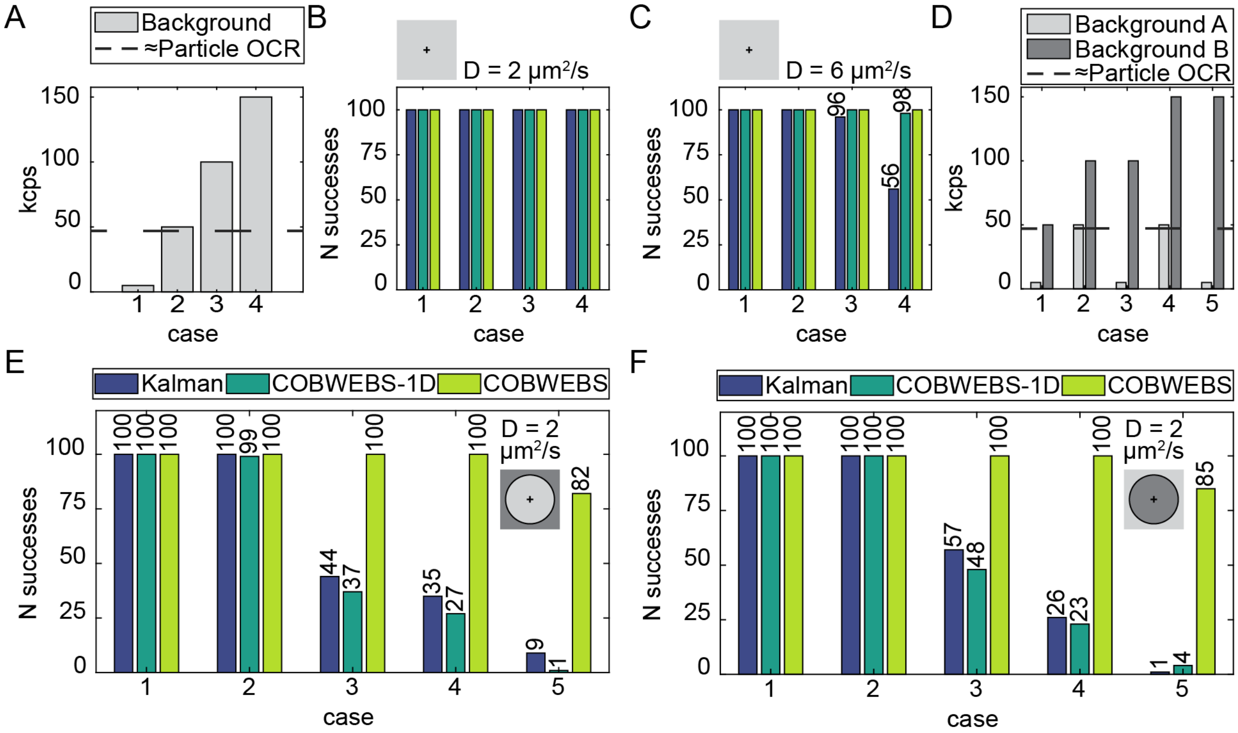

**Figure S10. Even and "dot" background cases. A.** Background levels for even background comparisons. **B.** Even background comparison trials for D = 2 µm²/s (100 trials, max duration 10 seconds, s = 516.64 kcps). **C.** Even background comparison trials for D = 6 µm²/s. **D.** Background intensities for dots explored in (E) and (F). **F.** Low to high dot pattern (100 trials, max duration 2 seconds, s = 516.64 kcps, D = 2 µm²/s). The background level is initially the lower of the two values from panel A then it increases to the higher value outside of 1.5 µm F-High to low dot pattern (100 trials, max duration 2 seconds, s = 516.64 kcps, D = 2 µm²/s). The background begins at the higher background level and then drops down to the lower value of a given case beyond 1.5 µm.

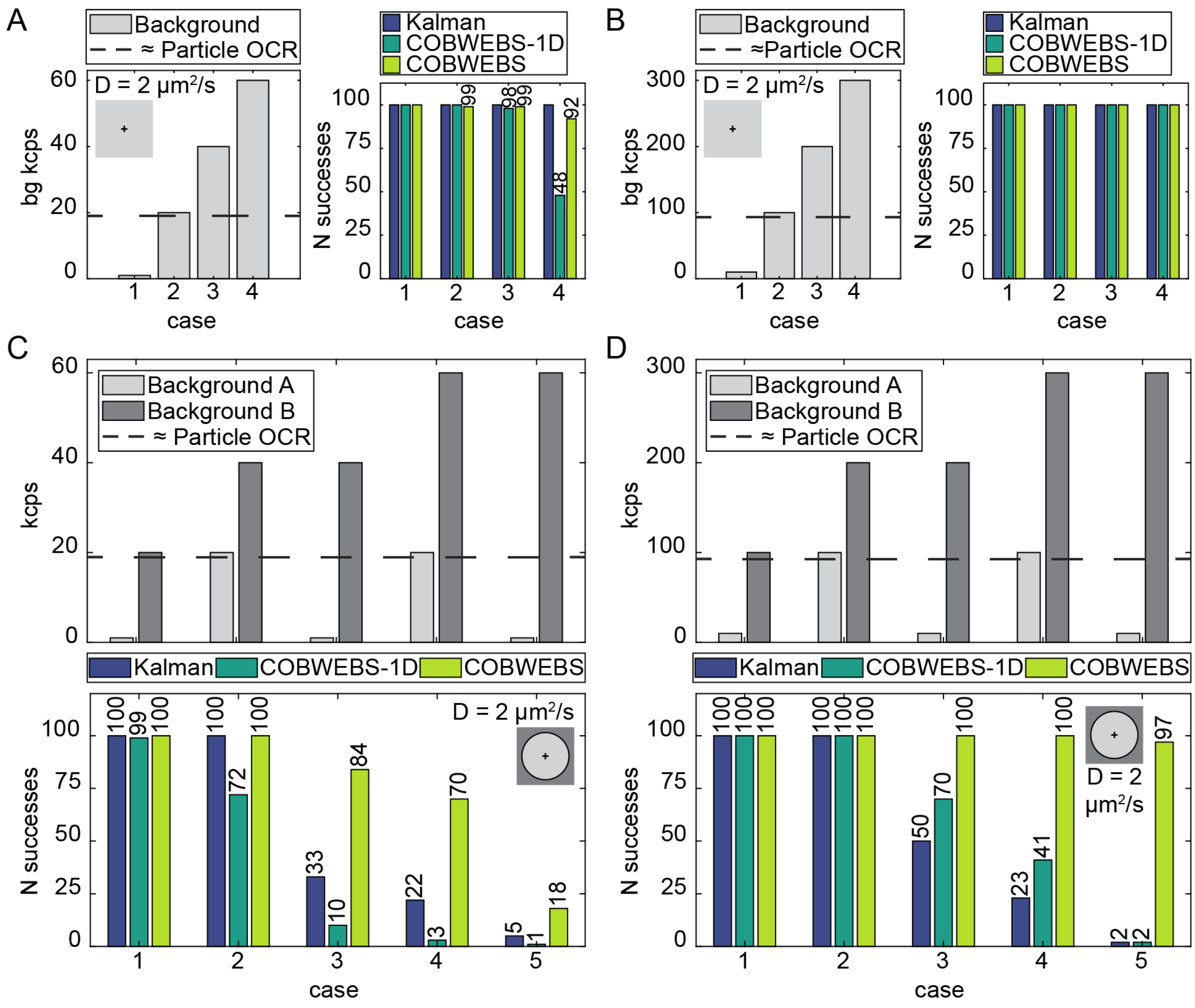

**Figure S11. Improved COBWEBS stability applies to both dimmer and brighter particles. A.** Even background comparison simulations for dim particle (100 trials, max duration 2 seconds, s = 206.656 kcps, D = 2 μm$^2$/s). **B.** Even background comparison simulations for a brighter particle, 2x the intensity of the VLP (100 trials, max duration 2 seconds, s = 1033.28 kcps, D = 2 μm$^2$/s). **C.** Top-background levels for dots. Bottom- N of full length trajectories for a dot going from low background (A) to higher background (B) at a distance from initial particle position of 1.5 μm. (100 trials, max duration 2 seconds, s = 206.656 kcps, D = 2 μm$^2$/s). **D.** Top-Background levels for dots. Bottom- N of full length trajectories for a dot going from low background (A) to higher background (B) at a distance from initial particle position of 1.5 μm. (100 trials, max duration 2 seconds, s = 1033.28 kcps, D = 2 μm$^2$/s).

#### SI Note 6: Impact of diffusion coefficient parameter estimate mismatch on tracking performance for additional particle intensities

##### 6.1 Additional particle intensities

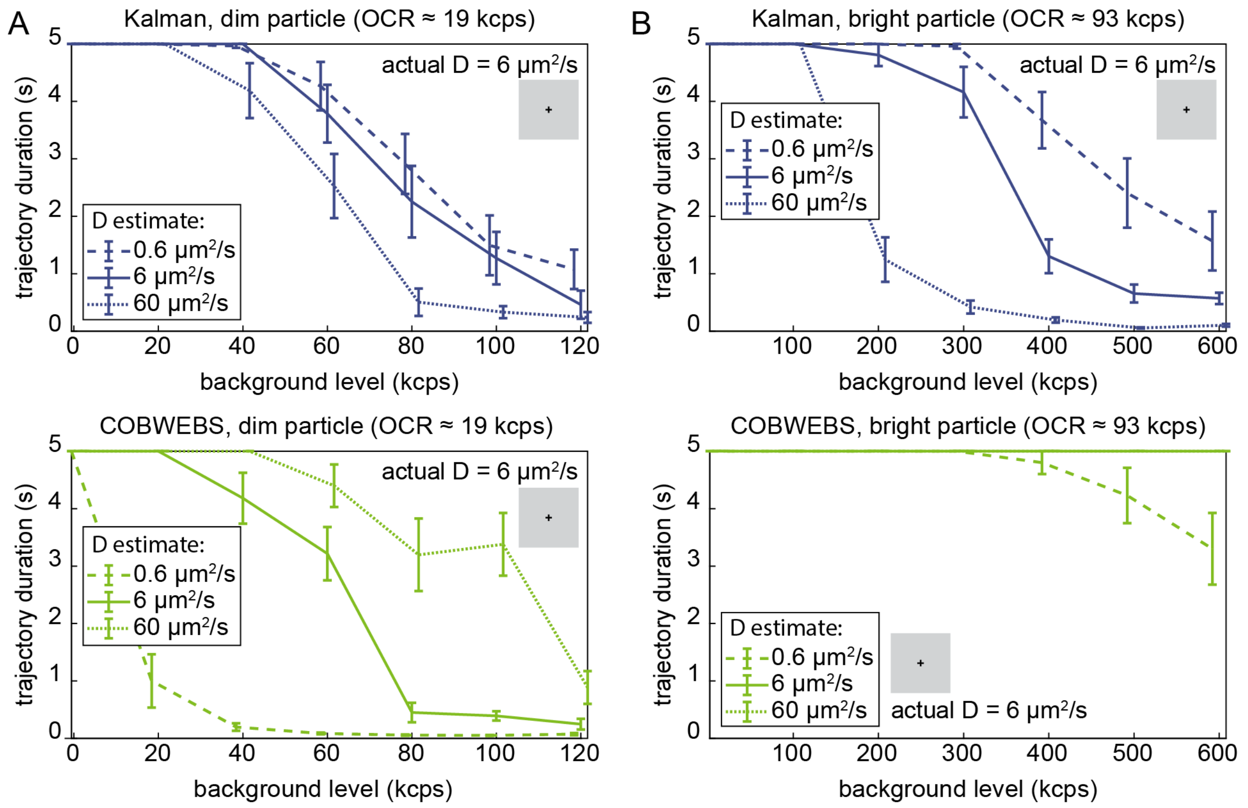

**Figure S12. Trajectory durations of both bright and dim particles with diffusion coefficient parameter mismatches in proportionally similar backgrounds.** 10 trials, max duration 5 seconds, s = 206.656 kcps or 1033.28 kcps, D actual = 6 μm²/s. Error bars are standard error of the mean.

Comparing dim particle tracking cases with perfect information, COBWEBS tracking is stable in backgrounds of up to 20 kcps while Kalman tracking appears to be stable up to 40 kcps (Figure S12A). Overestimation of the true diffusion coefficient for COBWEBS tracking by an order of magnitude improves trajectory durations making performance of COBWEBS tracking appear to meet or exceed Kalman tracking.

#### SI Note 7: Understanding dim particle tracking performance and the origin of the "trackable" versus "not-trackable" boundary

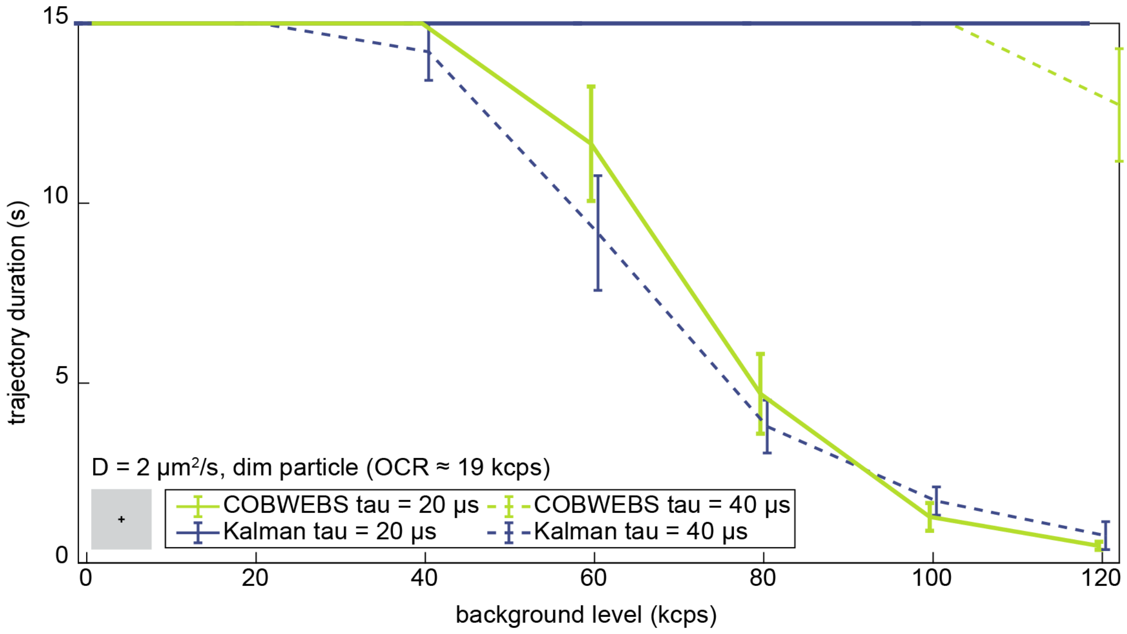

**Figure S13: Effect of increased bin times on trajectory duration for a dim particle.** 10 trials, max duration 15 seconds, s = 206.656 kcps, b = 1 to 120 kcps, D = 2 μm$^2$/s, tau: either 20 or 40 μs.

A potential solution to decreased COBWEBS tracking performance at low count rates is extending the bin time thus providing the algorithm with additional photons per update step. However, at the fast diffusion coefficient D = 6 μm$^2$/s increasing the bin time leads to failure of all tracking algorithms. For fast particles, increased motion blur cancels out any potential benefit (not shown). However, with a slightly slower particle diffusion of only 2 μm$^2$/s the hypothesized benefit to increased bin time for slowly diffusing particles in COBWEBS position estimation becomes evident as shown in Figure S13. The duration of these slower trajectories has been increased by a factor of 3 so that the tracked particles will cover the same mean displacement as the previous figures. Increasing the bin time from 20 μs to 40 μs improves COBWEBS trajectory durations to almost match those observed in simulated Kalman trajectories.

An additional experimental difficulty of tracking dim particles lies in identifying whether a particle is present in the laser scan area. Simulated trajectories were begun with particles already positioned within the laser scan area and terminated based on particle true positions. Once the particle's true position was more than 0.5 μm away from the edge of the scan area it was considered irrevocably escaped and the trajectory was considered failed. This gives the stage feedback time to catch back up to a particle which has ventured only slightly outside the scan radius and ensures that particles have fully escaped tracking prior to termination of trajectories. However, experimentally, one must use intensity to decide whether a particle is present in the laser scan and initiate or terminate stage feedback based on experimentally determined intensity thresholds. Due to noise in photon arrivals a Kalman filter is applied to the photon arrivals per bin which is then thresholded to identify particle presence (Figure S14A). The "thresholdability" is quantified by the overlap in filtered intensities between background levels alone and background + signal photons (Figure S14B). Extended over a range of particle and background intensities this metric corresponds nicely to the COBWEBS trackable vs not trackable boundary observed earlier and copied onto this figure. The Kalman trackable/not trackable boundary corresponds better to the signal to background ratio of the observed particle than this thresholdability metric.

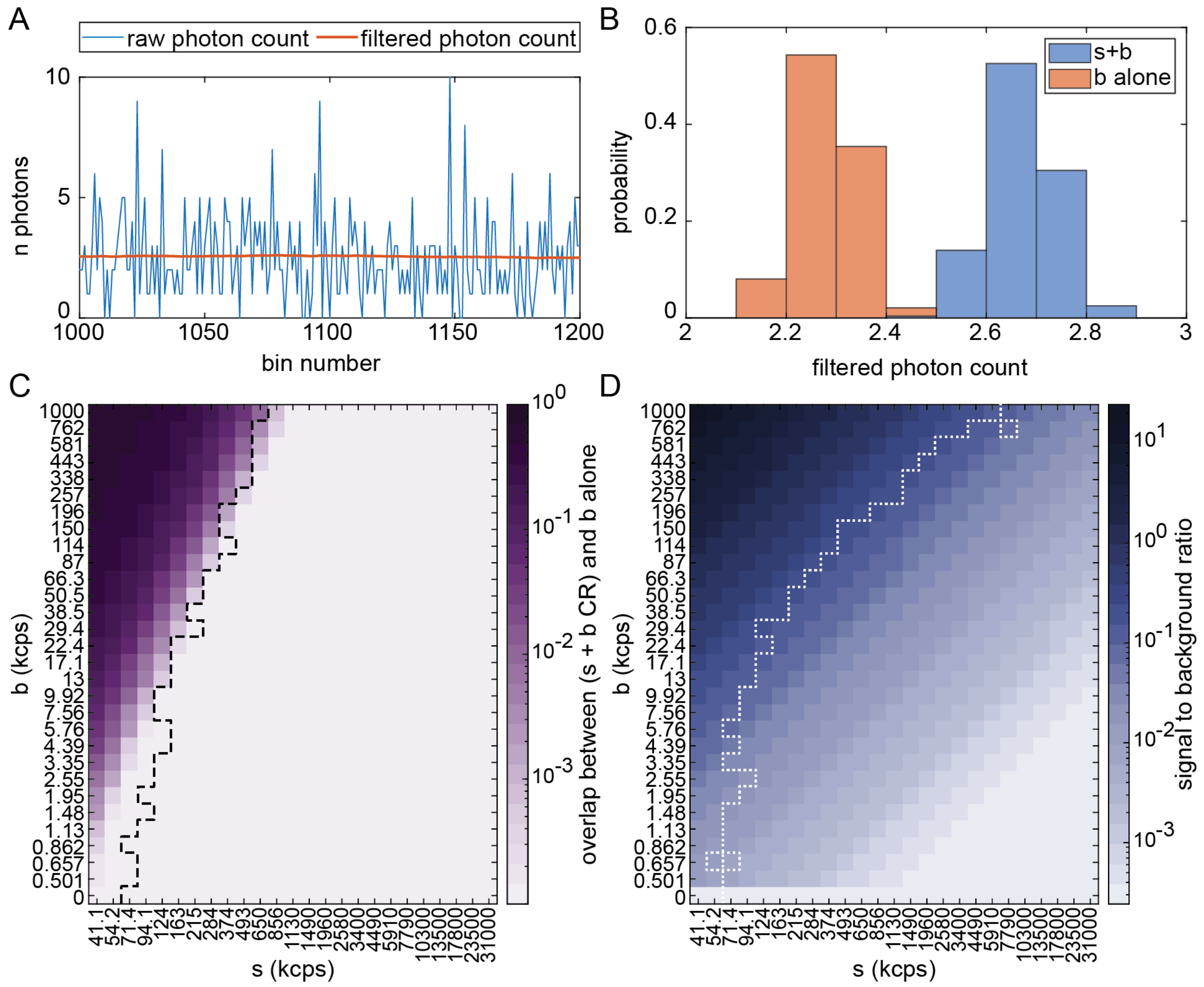

**Figure S14. Thresholdability and signal to background ratios. A.** Detected intensity for (s = 215335, b = 114062) both in raw photon arrivals and after Kalman filter with $k_i$ = 0.003. **B.** Histogram of filtered combined signal and background photons per bin and filtered background alone photon arrivals. Bin size displayed is wider than bin size used to calculate overlap integrals. **C.** Overlap integral between combined s and b photons and background only photon arrivals for range of particle and background intensities. Higher overlap indicates difficulty of distinguishing s and b photons. Trackable vs not boundary from Bayesian D = 6 $\mu m^2/s$ overlaid using black dashed line. **D.** Calculated signal to background ratio of particles tracked. Trackable vs not boundary from Kalman D = 6 $\mu m^2/s$ overlaid using white dotted line.

In summary, COBWEBS tracking is more challenging at low count rates. It may be possible to compensate for low count rates by extending bin times especially for slow moving particles. However, there may be cases where in theory for homogeneous backgrounds Kalman tracking is the more reliable tracking algorithm. In practice, Bayesian resilience to inhomogeneity may be the more relevant factor. Either way, experimental difficulty in thresholding whether or not particles are present in the laser scan may preclude tracking dim high background particles independent of algorithm selection.
